# Postmitotic separation enables selective niche retention of one daughter cell in intestinal crypts and is facilitated by interkinetic nuclear migration and basal tethering

**DOI:** 10.1101/142752

**Authors:** Thomas D. Carroll, Alistair J. Langlands, James M. Osborne, Ian P. Newton, Paul L. Appleton, Inke Näthke

## Abstract

Homeostasis of renewing tissues requires balanced proliferation, differentiation and movement. This is particullary important in the intestinal epithelium where lineage tracing suggests that stochastic differentiation choices are intricately coupled to position. To determine how position is achieved we followed proliferating cells in intestinal organoids and discovered that behaviour of mitotic sisters predicted long-term positioning. Normally, 70% of sisters remain neighbours while 30% lose contact separating after cytokinesis. Postmitotic placements predict differences in positions of sisters later: adjacent sisters reach similar positions; one separating sister remains close to its birthplace, the other moves upward. Computationally modelling crypt dynamics confirmed post-mitotic separation as a mechanism for placement of sisters into different niches. Separation depends on interkinetic nuclear migration, cell size, and asymmetric tethering by a basal process. These processes are altered when *Adenomatous polyposis coli* (*Apc*) is mutant and separation is lost. We conclude that post-mitotic placement enables stochastic niche exit and when defective, supports the clonal expansion of *Apc* mutant cells.

## INTRODUCTION

Fate choices of proliferating cells are critical for intestinal homeostasis. Lgr5(+) stem cells (SCs) in the intestinal crypt base must be regulated carefully to balance their maintenance with the production of transit-amplifying (TA) progenitors that can specialise. Similarly, exit of TA progenitors from their proliferative niche has to be regulated to produce the appropriate number of post-mitotic, differentiated cells. In the crypt, the position of cells relative to two niches, the stem cell and transit amplifying compartments, reflects their fate (Ritsma et al., 2014). Accordingly, stem and transit amplifying compartments differ in composition. The principal components of the intestinal SC niche are Paneth cells. Together with the surrounding mesenchyme, they provide Notch ligands, EGF and Wnts, which are critical for maintaining SCs and this creates a local Wnt gradient along the intestinal crypt axis (Sato et al., 2011). Displacement of stem cells from Paneth cell contact causes serial dilution of membrane-bound Wnts, contributing to loss of stemness (Farin et al., 2016). Neutral competition for niche access by the 12-16 SCs in the crypt base governs net contraction and expansion of clones, leading to mono-clonal crypts over time (Lopez-Garcia et al., 2010; Snippert et al., 2010). Stem cells near the border of the stem cell niche are more likely to enter the transit-amplifying compartment and lose stemness. Stem cells residing at or near the crypt base are more likely to retain stemness (Ritsma et al., 2014). Traversing the transit amplifying compartment is similarly accompanied by exposure to progressively less Wnt and other growth factors. Exit from this niche causes cell cycle exit. Such direct links between cell positioning and a graded niche signalling also operates in *Drosophilla* (Reilein et al., 2017). These observations suggest that in intestinal crypts, position, not the segregation of fate-determinants, regulates cell-fate.

Tissue homeostasis is perturbed in intestinal crypts mutant for key tumour suppressors such as *Adenomatous polyposis coli* (*Apc*), KRAS, p53 and SMAD4. These mutations provide cells with a selective advantage and increase their ability to colonise proliferative niches. Measuring the competitive advantage of cells carrying these mutations using sophisticated lineage tracing experiments demonstrated a competitive advantage over wild-type cells that allowed their preferential retention in the proliferative niche (Vermeulen et al., 2013, Song et al., 2014). The expansion of such mutant clones is thought to underpin field cancerisation, the preconditioning large tissue regions to neoplasia (Slaughter et al., 1953).

Our knowledge about cellular mechanisms that control cell positioning in the intestinal epithelium is limited, as is our understanding about how changes in such mechanisms can drive retention of mutant clones. Computational modelling suggests that the magnitude of the Wnt stimulus received at birth is a deciding factor for proliferative fate (Dunn et al., 2016). That suggests that decisions about cell position are set at birth. To test this hypothesis, we investigated daughter cell positioning along the crypt axis in 3D using intestinal organoids.

## RESULTS

We measured cell positioning during and after mitosis in intestinal organoids, a widely accepted physiological model of the intestinal epithelium (Sato et al., 2009). They contain epithelial domains that correspond to crypt-villus architecture *in vivo*, and contain a comparable cellular composition. Cell division (Figure 1A, S1 Figure) and polarity appear identical to those *in vivo* (Fatehullah et al., 2013), making organoids an ideal model system to understand the dynamic behaviour of the intestinal epithelium at temporal and spatial resolution impossible to achieve in tissue *in vivo*.

**Figure 1.**
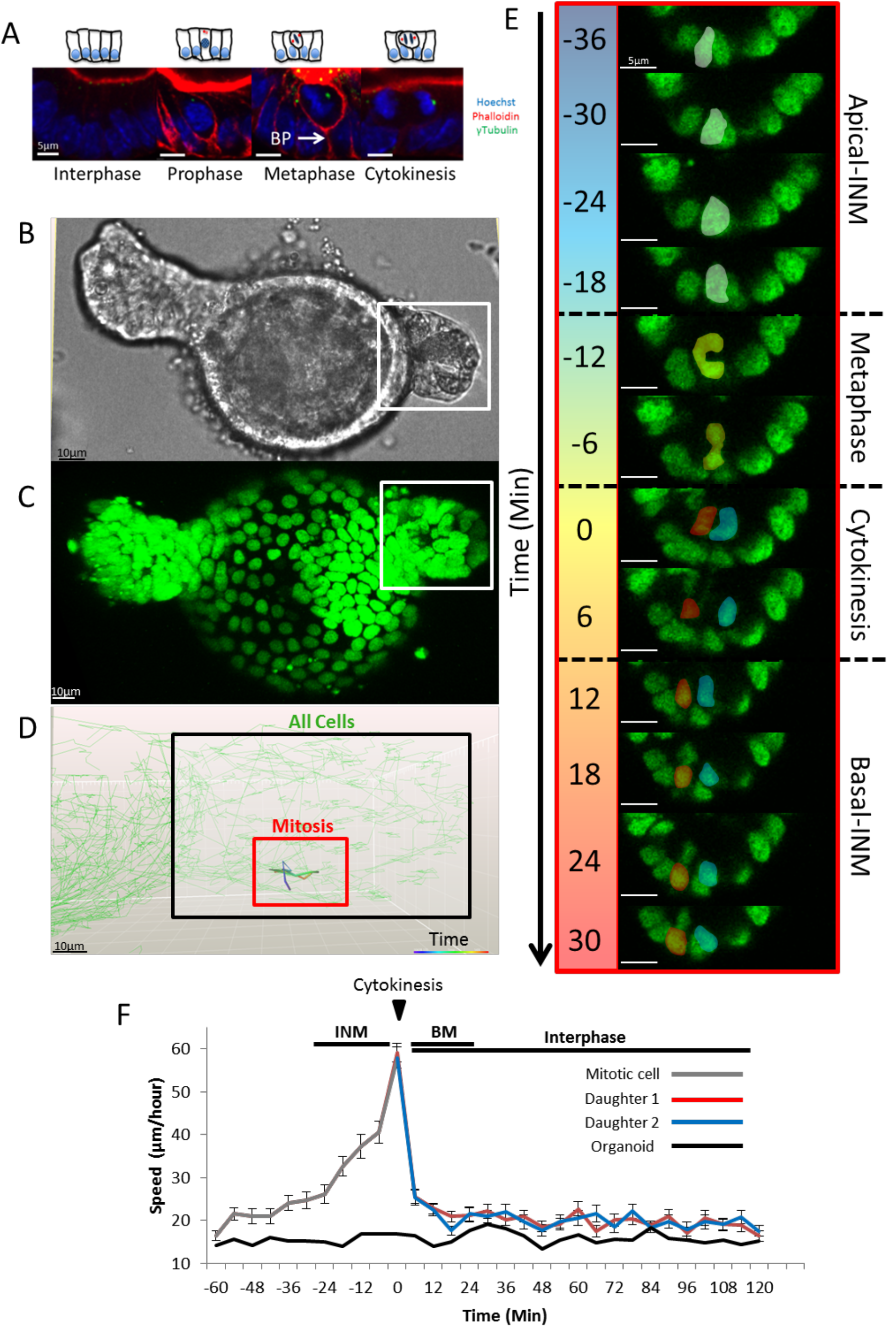
Dynamics of mitosis in H2B-GFP intestinal organoids. (**A**) Confocal sections of mitotic stages in intestinal organoids. Organoids were stained with Hoechst, phalloidin and γ-tubulin. (See also S1 Figure.) Representative bright-field (**B**) and fluorescent (**C**) images of a wild-type H2B-GFP intestinal organoid after 24 hours doxycycline treatment. (**D**) Manual tracking of a mitotic cell and its daughters. The track is colour-coded based on time (red box) and is overlaid onto the tracks of neighbouring cells (green), tracked automatically. In this example, tracks represent a time-lapse covering 66 minutes. (**E**) Dynamics of mitosis in intestinal organoids. Confocal sections (X-Y) of the mitotic cell highlighted in B. Prophase (white), metaphase (purple), cytokinesis (red) and daughter cell nuclei (blue and red) are shown. (**F**) Cell speed before, during, and after mitosis, measured for mother (grey line) and daughters (red/blue lines). Movement of the entire organoid was measured for reference (black line). The average speed was calculated for 60 mitotic cells from 3 different organoids. Data is displayed as mean +/− SEM. Time-points encompassing interkinetic nuclear migration (INM), cytokinesis, basal cell movement (BM) and interphase are highlighted.

### Interkinetic nuclear migration ( INM) operates in all intestinal epithelial cells and facilitates placement of mitotic sisters cells into different positions

Mitotic cells in the intestinal epithelium are easily distinguished (Figure 1A, S1 Figure). During interphase, nuclei are positioned basally. Upon entering mitosis, interkinetic nuclear migration (INM) causes nuclei to migrate apically towards the centrosome, similar to mitoses in the neuro-epithelium (Spear and Erickson, 2012). During this process, mitotic cells lose their columnar cell shape, become rounded and assume a position in the top half of the epithelial layer. Adjacent interphase cells expand into the basal space that is vacated by the migrating nuclei. Once INM is complete, spindles form and mitosis proceeds. After cytokinesis, newly formed cells move their nuclei basally and eventually assume columnar shape. As mitotic cells round up, their apical surface remains aligned with that of the epithelial layer and they remain attached to the basement membrane by a basal process. Centrosomes are located apically in interphase cells and align laterally with condensed chromosomes during metaphase. These mitotic stages are indistinguishable between tissue and organoids (S1 Figure).

### Dynamics of INM during mitosis

The distinct movement of pre- and post-mitotic nuclei in intestinal epithelium is similar to INM in other tissues, where it has been implicated in cell fate decisions (Spear et al., 2012). For instance, in the neuro-epithelium, INM facilitates differentiation by moving nuclei along apical-basal signalling gradients (Del Bene et al., 2008). In the developing foetal intestinal epithelium, INM has been implicated in the growth of epithelial girth (Grosse et al., 2011). The contribution of INM to intestinal homeostasis has never been examined. We examined how INM affects placement of mitotic sisters by tracking individual cells and their progeny during mitosis.

We directly monitored the position of mother and daughters during mitosis and after cytokinesis using live imaging of intestinal organoids expressing Histone2B-GFP (H2B-GFP). All nuclei in organoids derived from *H2B-GFP* mice robustly express GFP 24 hours after exposure to doxycycline allowing nuclear position to be used as a surrogate for cell position. (Figure 1B, C, S1 Movie) (Foudi et al., 2009). Measuring cell position in organoids required tracking cells in three-dimensional (3D) space. Techniques for accurately tracking cells in 3D are limited and we were unable to reliably track GFP(+) nuclei using automated methods. Therefore, daughter cell behaviour was recorded manually by tracking cells using Imaris (Bitplane) (Figure 1D).

Recordings revealed novel dynamic data about cell behaviour during mitosis. Mitosis lasted approximately 60 minutes. Prophase was characterised by nuclear condensation and INM, followed by rapid formation of the metaphase plate. After spindle alignment and cytokinesis, both daughters slowly migrate basally until their nuclei align with adjacent interphase cells (Figure 1E). During interphase, nuclei moved approximately 25μm/hour in crypts, which increased to 60μm/hour during INM. Speed during basal cell movement was comparable to that in interphase suggesting that INM is an active process and that basal movement is passive (Figure 1F). The unique arrangement of microtubule bundles above the nucleus in early mitosis suggests that INM involves microtubules (S1 Figure).

### Daughter cells either remain adjacent or are separated from one another after mitosis

Tracking mitotic cells revealed two distinct outcomes for mitotic sisters. They either remain adjacent (6.0 +/−1.2μm apart) and become neighbours (Figure 2A, S2 Movie), or they separate (12.9 +/− 2.8μm apart) and exchange neighbours (Figure 2B, S3 Movie). Rendering mitoses in 4D confirmed their separation by a neighbouring cell (Figure 2C, S4 Movie). Importantly, we observed similar mitoses *in vivo* with one sister positioned significantly displaced from the other by neighbouring cells (Figure 2D). This data suggests that postmitotic separation occurs in native tissue and in organoids.

**Figure 2.**
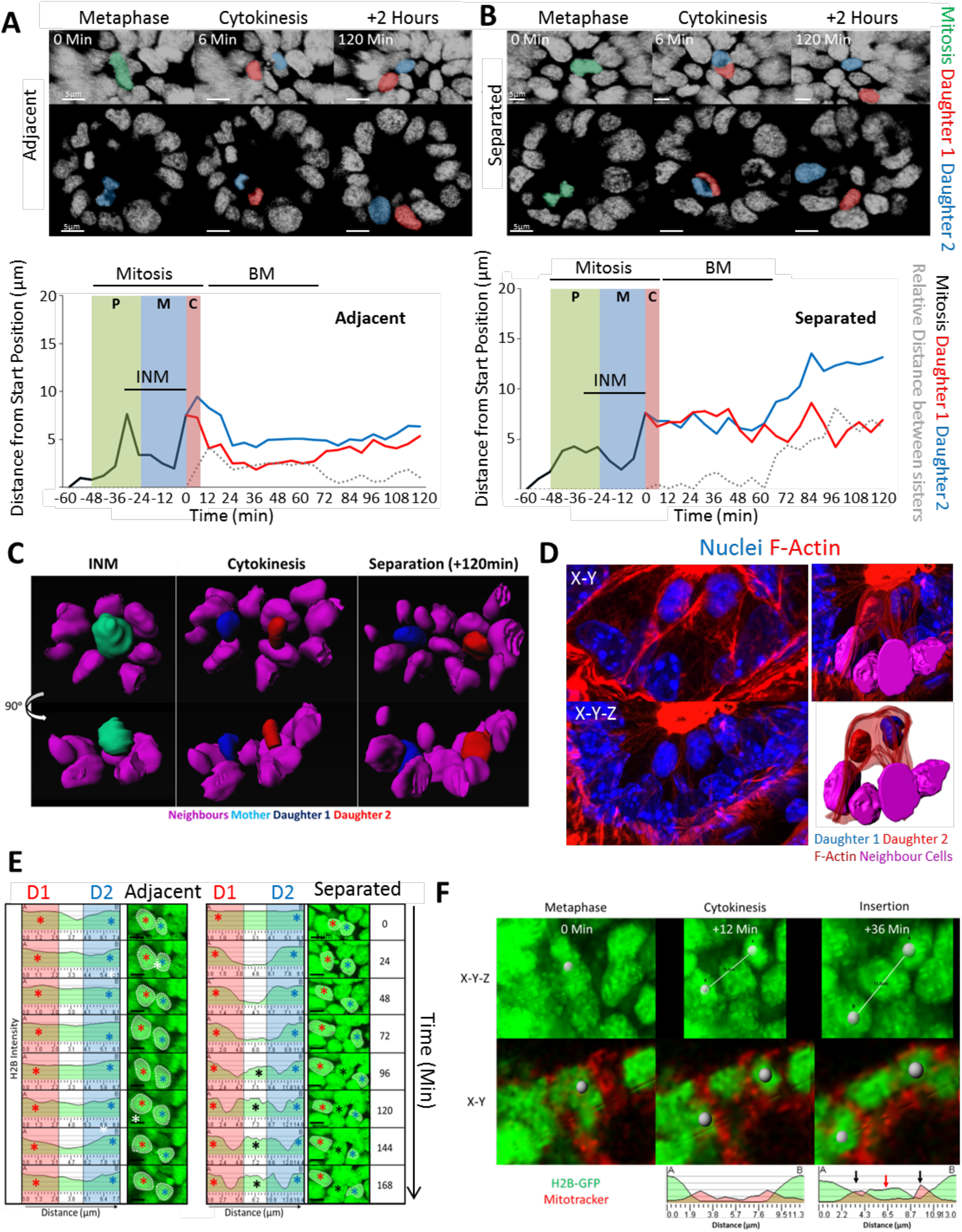
Post-mitotic separation of daughter cells. Mitotic cells were tracked manually for 60 minutes prior to cytokinesis and daughters for a further 120 minutes. Two types of mitotic types were revealed: (**A**) Daughter cells positioned adjacent or (**B**) which separated after mitosis. Displayed are 3D projections (top panels) and 2D sections through an organoid branch. Metaphase (green) and daughters (red/blue) are highlighted. Representative tracks show the distance of the mitotic mother (black line) and daughters (red/blue lines) from the original starting position. (P)rophase, (M)etaphase, (C)ytokinesis, interkinetic nuclear migration (INM), and basal cell movement (BM) are indicated. Distances between adjacently placed daughters (grey dashed line) are ≤ 1 nuclear width (6μm) whereas distances between separating daughters is greater. (**C**) 3D rendering of neighbouring nuclei (purple), mother (cyan) and daughters (red/blue) of a post-mitotic separation event. Displayed are rotated views of cells and their direct neighbours at time-points encompassing INM, cytokinesis and after separation (120 minutes after cytokinesis). (**D**) Daughter separation occurs *in vivo*. Representative image of daughters at a crypt base. Samples were stained with Hoechst (blue) and phalloidin (red). Highlighted are two prospective daughters (white stars) displayed in X-Y and X-Y-Z views (left panels). Surface rendering (right panels) highlights cell-cell boundaries and neighbouring cell nuclei. (**E**) H2B-GFP line-intensity profiles were created along a line connecting the centres of sister nuclei at indicated times after cytokinesis (=Time 0). Reference images (3D projections) are displayed. Please note that the scaling for the x-axis, indicating distance, changes in the right-hand panel to accommodate the increased space between the separating daughters. (**F**) Individual frames of an H2B-GFP organoid stained with Mitotracker highlighting a mitotic cell whose daughters separate shortly after mitosis. Time points reflecting metaphase, cytokinesis and after return of daughters to their interphase position (reinsertion) are shown. A line-intensity profile was generated between marked daughters (A and B) during cytokinesis and after ‘insertion’. After reinsertion, a discrete H2B-GFP peak was detected that correspond to the neighbouring cell that displaces the two daughters (red arrow). The neighbouring cell has two distinct Mitotracker peaks on either side of its H2B-GFP signal (black arrows).

To determine when mitotic sisters separate, we measured when neighbouring cells first appeared between them. Specifically, we measured the H2B-intensity across the line connecting the centre of sister nuclei to visualise nuclear boundaries (Figure 2E). For adjacent sisters, the line-intensity profile was unchanged over time indicating that the two nuclei remained in close proximity. In post-mitotic separations, an additional peak appeared between the peaks representing each sister, indicating the insertion of a neighbouring cell between them. Insertion of neighbours occurred 72-120 minutes after cytokinesis, indicating that displacement occurred during basal cell movement (Figure 2E). Live-imaging of the mitochondrial network using Mitotracker clearly showed distinct cells between mitotic sisters, further confirming their physical separation (Figure 2F, S5 Movie).

We found other situations also favoured separation. Separation could be facilitated by the movement of daughters of other mitoses in the immediate vicinity (S2 Figure, S6 Movie). Furthermore, separation was favoured when mitoses occurred next to Paneth cells. Paneth cells are more adherent and stiffer (Langlands et al., 2016) and this could force one daughter cell out of the way (S2 Figure; S7 Movie)

### *Apc* mutation alters placement of daughter cells

APC is required for normal intestinal homeostasis and mutations in *Apc* are common to most tumours in the colon (Fearnhead et al., 2001). The APC protein functions as a scaffold in Wnt signalling (McCartney and Näthke, 2008). It contributes to spindle orientation (Yamashita et al., 2003, Quyn et al., 2010), and cell migration along the crypt-villus axis (Nelson and Nathke, 2013). Lineage tracing and computational modelling has demonstrated that *Apc* mutations increase the retention of cells in intestinal crypts (Vermeulen et al., 2013, Song et al., 2014). To determine if changes in the positioning of mitotic sisters could explain these observations we isolated organoids derived from *Apc* heterozygous mice (*Apc^Min/+^*). These organoids are initially indistinguishable from wild-type organoids (Fatehullah et al., 2013) but transform into spherical, cyst-like structures (Figure 3A) containing cells that have undergone loss of heterozygosity (LOH) (*Apc^Min/Min^*) (Germann et al., 2014). Mitoses appeared normal in *Apc^Min/+^* organoids, however, in *Apc^Min/Min^* organoids, abnormal mitoses with multipolar spindles and mitotic slippage were frequently observed (S3 Figure), similar to cultured cells which lack APC (Dikovskaya et al., 2007). We compared the incidence of the two types of cell placements in wild-type and *Apc^Min/+^* mice and in *Apc^Min/Min^* organoids (S1 Movie).

**Figure 3.**
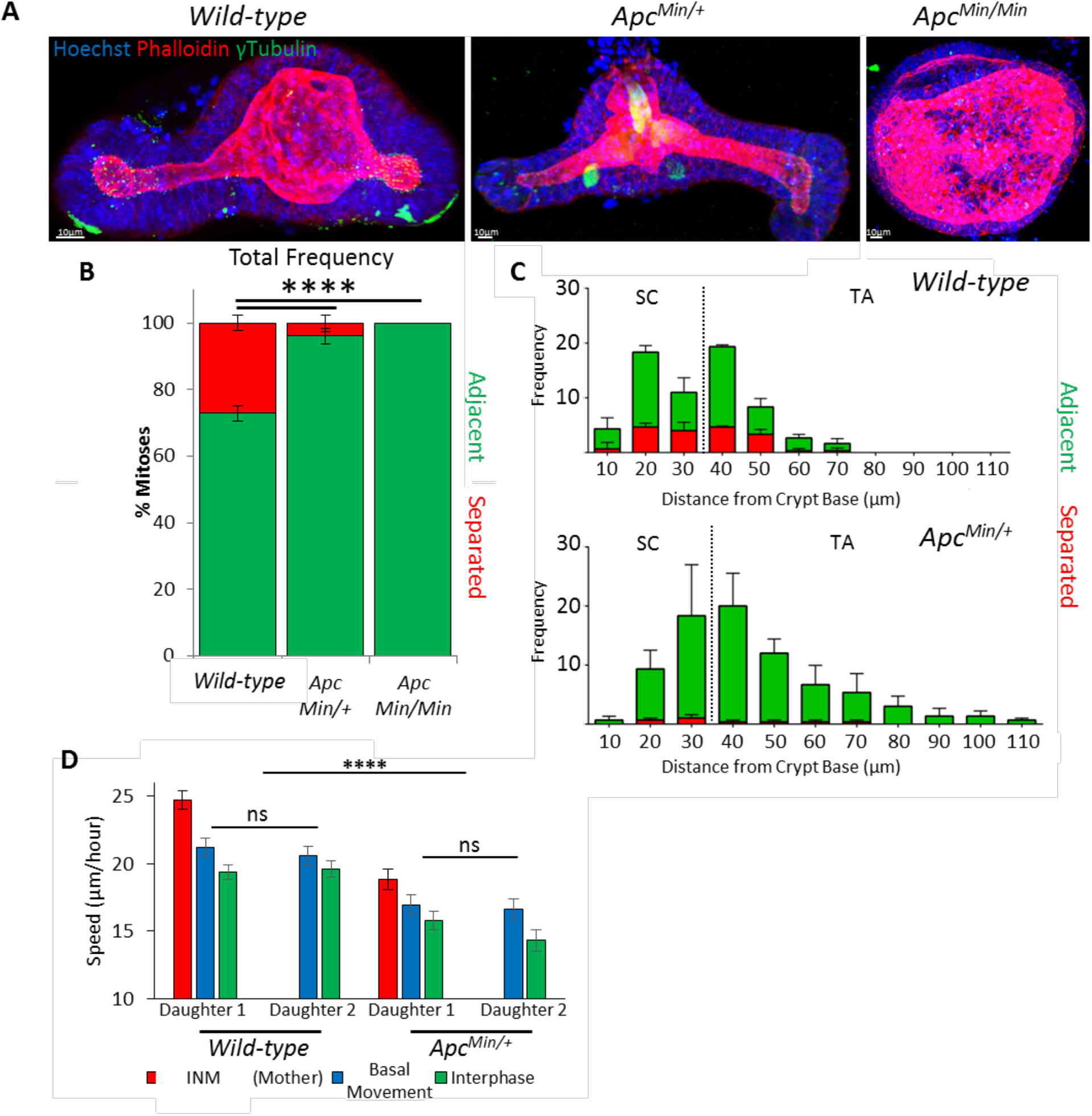
*Apc* mutant daughters separate less frequently. (**A**) 3D projections of fixed organoids produced from small-intestinal crypts of wild-type and *Apc^Min/+^* mice stained with Hoechst (blue), phalloidin (red) and γ-tubulin (green). *Apc^Min/+^* organoids from cells that have undergone LOH form cysts (*Apc^Min/Min^*). (See also S3 Figure.) (**B**) Types of mitotic daughter placement were scored in organoids (wild-type N=6, 491 mitoses; *Apc^Min/+^*, N=3, 227 mitoses, *Apc^Min/Min^*, N=7, 34 mitoses; T-test). Relative frequency of each type of mitosis was determined per organoid and averaged for replicate organoids. There was a significant difference between the number of adjacent and separating daughters between wild-type, *Apc^Min/+^* and *Apc^Mm/Min^* organoids (T-test, p<0.0001). (**C**) Mitotic cell position was determined relative to the crypt base for wild-type and *Apc^Min/+^* organoids. The frequency of each mitosis type along the crypt-villus axis was measured for 3 organoids. The stem cell (SC) and transit amplifying (TA) compartments are marked as defined by the average position of Lgr5-GFP(+) cells (see S5 Figure). Data is displayed as mean +/− SEM. (**D**) Nuclear speed was calculated in wild-type and *Apc^Min/+^* organoids (N=3 organoids, 20 cells) showing average speed for interkinetic nuclear migration (INM), basal cell movement and interphase. Data is displayed as mean +/SEM. There was a significant difference between the speed of cells in wild-type and *Apc^Min/+^* organoids (T-test, p<0.0001).

In wild-type epithelium, ca. 30% of daughter cells separated whilst ca. 70% remained adjacent (Figure 3B). Separation was mainly associated with movement of neighbouring interphase cells during basal cell movement (75.1% +/− 14.8% of cases). Separation by surrounding mitotic progeny was less common (29.5% +/−21.6% of cases). The frequency of the two mitotic types was equal in the stem and transit amplifying compartments, suggesting that mitotic outcome is independent of cell position and type and can occur in any cycling cell that undergoes INM (Figure 3C). To further confirm that mitotic separation is not specific to stem cells, we measured mitotic outcome in organoids treated with the GSK-3β inhibitor, Chir99021, and the HDAC inhibitor, valproic acid, which increases the number of Lgr5(+) stem cells in the crypt (Yin et al., 2014). Treatment with Chir99021 and valproic acid did not significantly change post-mitotic separation of sisters (S4 Figure), suggesting that the occurrence of post-mitotic separation is similar in all dividing cells along the crypt axis.

In *Apc^Min/+^* organoids there was a significant reduction in the frequency of postmitotic separations. Sisters never separated in *Apc^Min/Min^* organoids (Figure 3B). This suggests that *Apc* mutant sisters are more likely to remain adjacent after division. There was also a significant overall reduction in cell movement between wild-type and *Apc^Min/+^* epithelial cells, including nuclear speed during INM (Figure 3D), suggesting that cells remain adjacent because of reduced cell movement. Loss of post-mitotic separation was also induced by long-term treatment of organoids with high concentrations of Chir99021. This treatment caused organoids to grow as cysts, similar to *Apc^Min/Min^* organoids (S5 Figure). This suggests that hyperactive Wnt signalling induced either by *Apc* mutation or by GSK-3β inhibition can alter the frequency of post-mitotic separation, although it is possible that this is an indirect consequence of the changes in cells size and shape (see below).

### Post-mitotic separation of daughter cells directs niche exit

The two types of placements of mitotic sisters we discovered led to the hypothesis that post-mitotic separation allows differential exit of sisters from proliferative compartments. For instance, separation of stem cell daughters may increase the probability for one daughter to remain in the stem cell niche compared to the other. Similarly, in the transit-amplifying compartment, post-mitotic separation could make it more likely for one daughter to remain in the proliferative compartment and the other to exit and terminally differentiate. To test this idea, we measured the distance of mitotic sisters from their starting positions and from each other after their birth. Shortly after cytokinesis, after both daughters had assumed their interphase position, regardless of mitosis type, one sister always remained near its starting position, whereas the other moved upward (Figure 4A, B). At later times (up to 35 hours after mitosis), differences between sisters were accentuated. If sisters had separated, one always remained close to its starting position while the other was displaced significantly upwards. In contrast, adjacently placed sisters were both displaced upwards (Figure 4A, B). Thus the initial difference in distance between sisters in the two types of mitoses was amplified over time, consistent with the idea that different placement of mitotic sisters can produce different outcomes for cell positioning.

**Figure 4.**
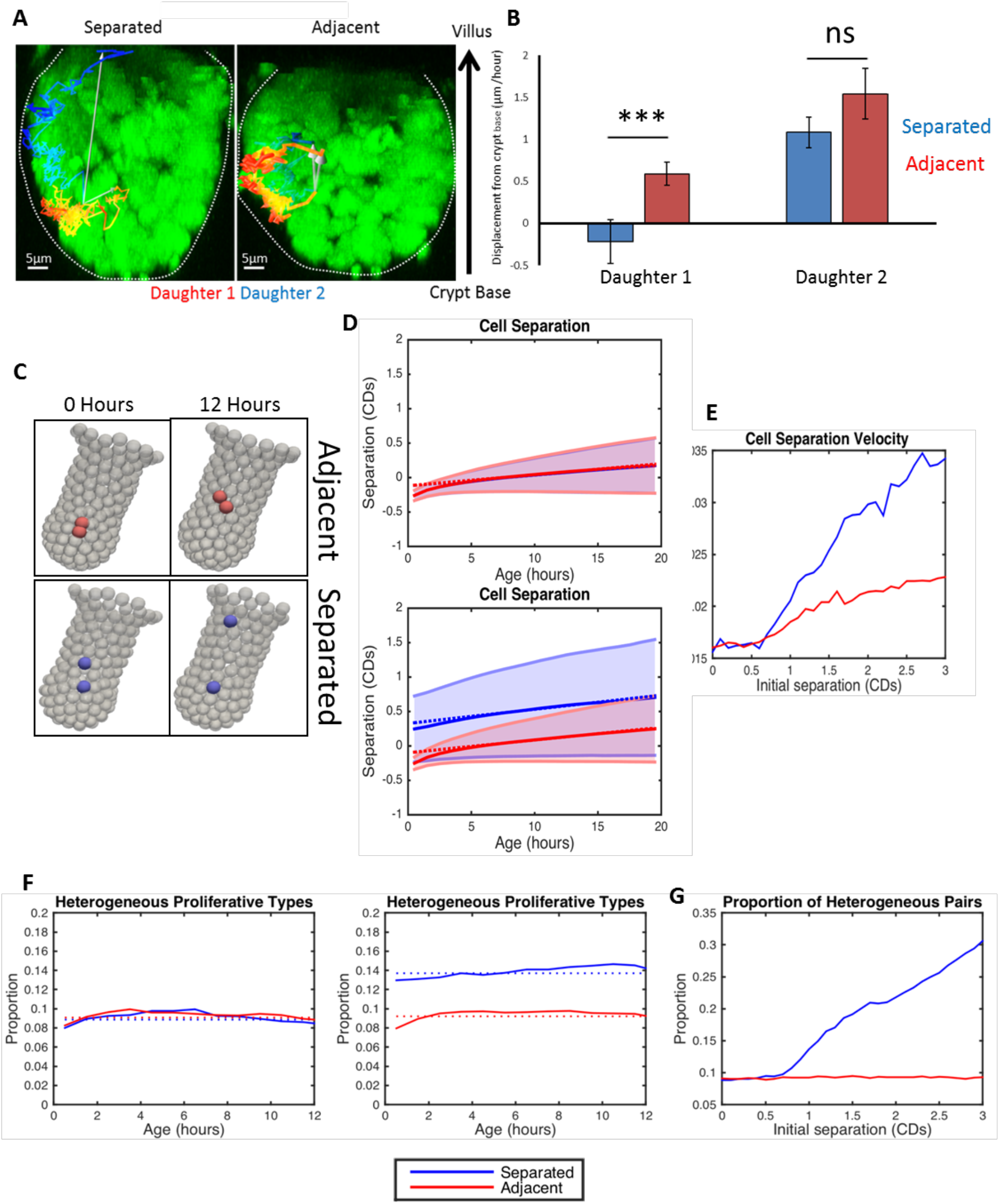
Post-mitotic separation promotes both niche retention and exit. (**A**) Representative images show examples of the long-term behaviour of daughter cells following separate or adjacent placement. An overlay of the track of daughters (Daughter 1, red; Daughter 2, blue) reveals the total displacement over the time course (white arrow). (**B**) Daughter cell position from the crypt base was measured after reinsertion into the epithelium (^~^2 hours) and at the final position able to be recorded (5-35 hours). Displacement (movement of each daughter from the crypt base over time) was calculated for each daughter pair. Daughter 1 was defined as the daughter closest to the crypt base. Values were calculated for separating (Apart, N = 28) or adjacent (Together, N = 84; T-test, p<0.001) sisters. (**C**) Simulation results: representative images showing daughters initially placed adjacent to each other (red) and placed apart by 1 cell diameter (blue). Snapshots shown represent these situations immediately after mitosis and 12 hours later (right hand panels). (**D**). Simulation results show the distribution of cell separation as a function of time since birth. Results are shown for a homogenous population (left) of cell divisions where daughter cells are placed adjacent to each other (i.e. *S* = 0 for all divisions), and for a heterogeneous population (right) of cell divisions where *S* = 0 for two thirds of divisions (red) and *S* = 1 for the remaining third (blue). The mean separation (solid line) and standard deviation (shaded region) is displayed. Linear fits to the distribution (from 5-15 hours) are represented by dotted lines. (**E**) Simulation results for the effect of initial cell placement on separation velocity. For each separation, a heterogeneous population of divisions (two thirds with *S* = 0 and a third with *S* ≠ 0) is simulated and the corresponding separations (as shown in (D)) are calculated, and the distribution of values recorded. The separation velocity is calculated by taking the gradient of the linear fit to the mean of this distribution for both populations of divisions (adjacent in red and separated in blue). Cells placed initially further apart will separate more quickly than those placed together. (**F**) Simulation results for the proportion of cell divisions that produce cells in different niches (i.e one cell remains in the proliferative compartment while the other leaves) for simulations shown in (D). Results are shown for a homogenous population (left) of cell divisions where all daughter cells are placed adjacent to each other (i.e. *S* = 0 for all divisions) and for a heterogeneous population (right) of cell divisions where *S* = 0 for two thirds of divisions (red) and *S* = 1 for the remaining third (blue). Constant fits to these distributions (using data for all ages) are denoted by dotted lines. (**G**) Simulation results showing how the average proportion of cell pairs with different positions (using the constant fit from (F)) depends on the initial separation. The same simulations were used as in (E). Increasing separation leads to a larger proportion of cell pairs in different positions.

To provide additional evidence for this idea we used a previously established computational 3D model of intestinal crypts (Dunn et al., 2016) and asked whether postmitotic separation could promote heterogeneous position/fate. To compare modelling results to our experimental data, simulations were performed with daughters placed adjacent to each other (as in previous computational models) or separated by a factor larger than a typical cell diameter. Simulations were performed using parameters derived from the primary data (materials and methods). These simulations confirmed that post-mitotic separation often led to one daughter being retained close to its point of birth whilst the other displaced upward (Figure 4C). There was a significant difference between the separation velocities between the two mitotic subtypes, indicating that daughters that initially separated moved apart much faster than those born adjacent (Figure 4D, E).

To test whether post-mitotic separation influences the number of heterogeneous cell pairs, we imposed a crypt-specific boundary separating a proliferative region from a non-proliferative region. Heterogeneous pairs are produced when one daughter is retained in the proliferative boundary and the other exits. Consistent with our experimental results, simulations showed that separation led to more heterogeneous pairs than adjacent placements (Figure 4F). The same results were produced for thresholds representing the TA/Differentiated SC/TA boundaries, similar to other reports (Vermeulen et al., 2013, Song et al., 2014). A greater separation distance at birth led to a higher number of heterogeneous pairs (Figure 4G). Together, these data suggest that post-mitotic separation could enhance divergent daughter fate by promoting the exit of one daughter from a niche whilst allowing the other to remain.

### Mechanisms for post-mitotic separation

A number of mechanisms may be involved in post-mitotic separation. For instance, spindle orientation could direct placement of sisters, supported by the different types of spindle alignment we previously discovered in intestinal tissue (Quyn et al., 2010). Position and location of mitotic sisters is likely affected by spindle orientation. To understand how mitotic placement related to spindle orientation, we measured spindle alignment in organoids. Consistent with previous data in whole tissue, we observed spindle orientation bias in the stem cell compartment of wild-type organoids where cells more readily oriented their spindles perpendicularly to the apical surface. There were no perpendicularly oriented spindles in the stem cell compartment in *Apc^Min/+^* organoids (S3 Figure). This is consistent with the idea that separating sisters can result from mitoses with perpendicularly aligned mitotic spindles. However, perpendicular spindle alignment was less frequent than separating sisters, indicating that additional processes are involved and that spindle orientation is not a reliable measure of post-mitotic separation.

### Basal tethering of daughter cells contributes to post-mitotic separation and is altered in *Apc* mutant organoids

Another mechanism that may affect placement of daughters involves the basal process that tethers mitotic cells to the basement membrane (Fleming et al., 2007). This process is formed during INM and persists throughout metaphase. The basal process is rich in F-Actin and is tethered at the basement membrane by β4-Integrin (Figure 5A). Tethering of daughter cells after cytokinesis and during basal cell movement provides a direct means to guide daughters. Asymmetric tethering of mitotic cells has been shown to coincide with the segregation of planar cell polarity markers in the colonic epithelium (Bellis et al., 2012).

**Figure 5.**
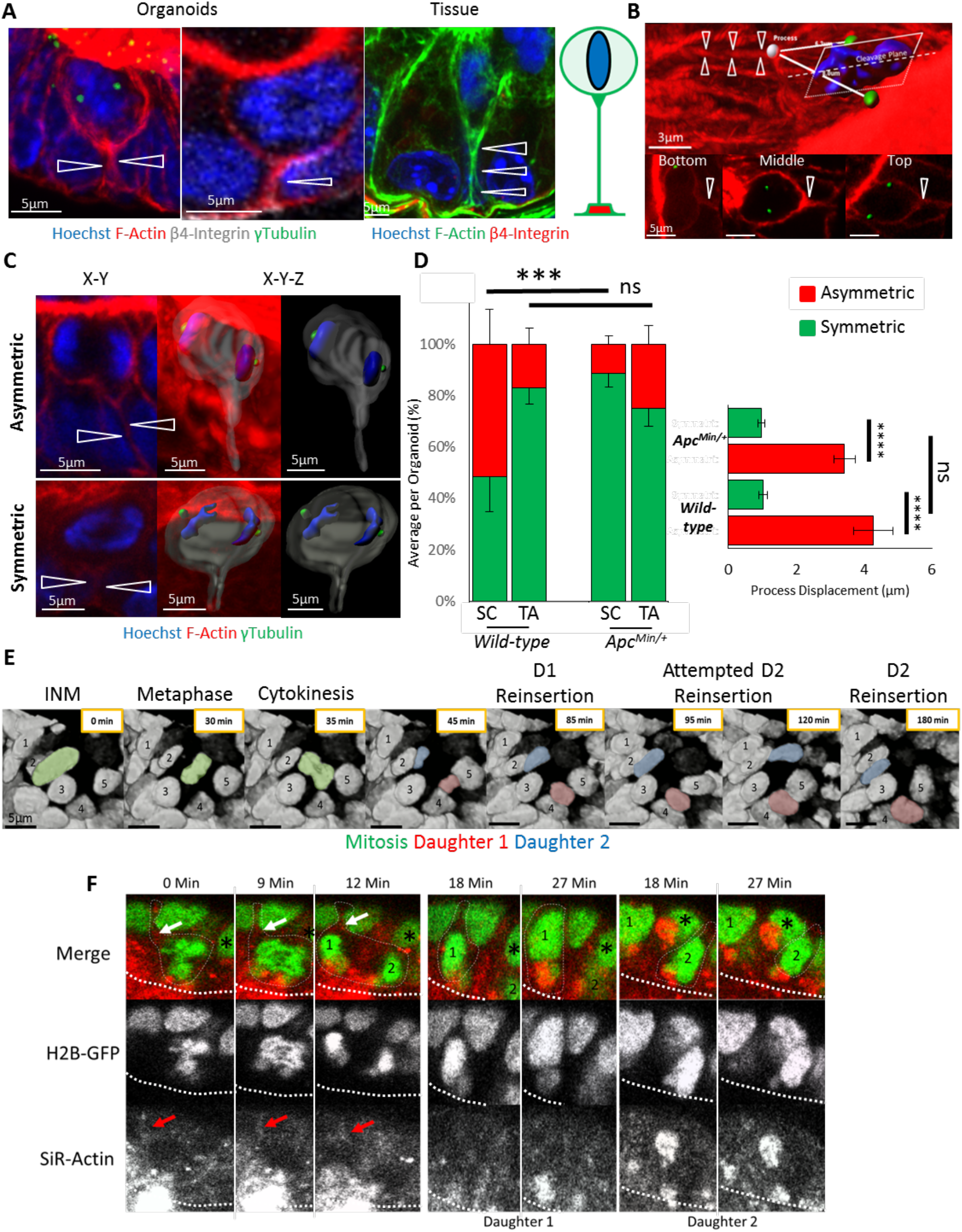
Basal tethering of mitotic cells is altered in Apc mutant epithelia. (**A**) 3D projections of mitotic cells in organoids and whole tissue reveal the basal process (white arrows). Samples were stained with Hoechst (blue), phalloidin (red), γ-tubulin (green) and β4-Integrin (white). A schematic of the basal process is shown. (**B**) 3D projection of a mitotic cell aligned in metaphase shows the position of its basal process (white arrow), centrosomes (green), nucleus (blue), and mitotic cleavage plane. Symmetric or asymmetric process inheritance was scored based on its placement relative to each centrosome. Accordingly, basal processes could be localised to the cell closer to the crypt base (‘bottom’), equidistant from each centrosome (‘middle’), or furthest from the crypt base (‘top’). In the displayed X-Y sections, views were orientated with the crypt base towards the bottom in each image. (**C**) Process inheritance was scored by visual inspection of the position of basal processes relative to the mitotic cell in 3D. Two examples of mitotic cells are shown, one with asymmetric and one with symmetric process placement (see also S3 Figure). 3D surface rendering shows the position of the basal process. (**D**) Process segregation was scored by measuring the distance between the attachment point of the basal process and the centrosome of each prospective daughter in the stem cell- and transit-amplifying compartments of wild-type (N=12 organoids, N = 68 mitoses) and Apc^Min/+^ (N=20 organoids, N = 61 mitoses) organoids. Frequencies are displayed as the average percentage of each outcome per organoid. The frequency is displayed as a percentage of the mitotic events in each compartment. There was a significant reduction in the number of asymmetrically localised basal processes in the stem cell compartment in Apc^Min/+^ organoids compared to the stem cell compartment in wild-type organoids (T-test ***p<0.001). Processes were scored manually and defined as asymmetric if significantly displaced from the cleavage furrow. To confirm manual scoring, process displacement was calculated for all scored asymmetric and symmetric processes. Displacement was defined as the difference between the distances from the process to each centrosome (right hand panel). Data is displayed as mean +/− SEM. Process displacement in mitoses scored as asymmetric was significantly more common than in symmetric mitoses (T-Test ****p<0.0001). (**E**) Individual frames of a time-lapse movie reveal the repeated attempt of one daughter (red) to assume the original position of the mother (green) while the other daughter (blue) moves on. (**F**) Individual frames of a time-lapse of H2B-GFP organoids stained with SiR-Actin show a cell whose daughters undergo post-mitotic separation. Displayed are time points encompassing metaphase (0 min), cytokinesis (9-12 min) and the two daughters during reinsertion (18-27 min), when they become separated by a neighbour (black stars). An asymmetric process (white/red arrows) is located closer to daughter 1 on one side of the putative cleavage furrow. The apical surface is denoted by the thick dashed line.

To determine whether asymmetric tethering of sisters operated in organoids and contributed to their placement, we measured the position of basal processes relative to prospective daughters. We distinguished whether the process was positioned symmetrically or asymmetrically. Processes attached close to the cleavage plane, equidistant to both centrosomes, were classified as symmetric. Those attached closer to one centrosome were classified as asymmetric. For asymmetrically placed processes, we also measured their position relative to the crypt base, i.e. whether the mitotic sister they were connected to was closer to the bottom or top of the crypt (Figure 5B, C). The basal process in all separating mitoses was significantly more displaced from the cleavage furrow than in adjacent mitoses (Figure 5D). Accordingly, symmetrical processes predicted equal tethering of daughters and adjacent cell placement, whereas asymmetric processes predicted daughter separation (Figure 5C).

The proportion of symmetrically and asymmetrically placed basal processes differed between the stem cell and transit-amplifying compartments. In the latter, the proportion of asymmetric basal processes was ca. 30%, similar to the proportion of separating mitotic daughters. However, in the stem cell compartment, this number increased to 50% (Figure 5D). In both regions, asymmetrically positioned processes tended to localise to the daughter cell closer to the crypt base, predicting that the untethered daughter was most likely to be displaced upwards. Asymmetrical basal process placement was a feature of mitotic cells with perpendicularly aligned spindles, suggesting that spindle orientation and basal process placement are linked (S3 Figure). Live imaging suggested that the basal process guides basal cell movement. The tethered daughter migrated basally to assume the interphase position of the mother, whilst the untethered daughter moved freely and allowed sister separation. This was particularly obvious when a daughter required multiple attempts to reintegrate into the epithelium (Figure 5E; S8 Movie). In *Apc^Min/+^* intestinal organoids fewer processes were placed asymmetrically, consistent with the significantly reduced frequency of separating sisters in *Apc^Min/+^* organoids (Figure 5D). We propose that asymmetric processes facilitate the displacement of one daughter cell from the niche by allowing it to separate from its sister, rather than simply aiding in their retention (Bellis et al., 2012). To provide evidence for this hypothesis, we performed live imaging of H2B-GFP organoids treated with SiR-Actin (Lukinavicius, G. et al., 2014). As expected, daughter separation correlated with asymmetric segregation of the basal process (Figure 5F, S9 Movie).

### *Apc* loss and hyperactive Wnt signalling restrict separation of sisters by inhibiting INM and changing cell size and morphology

We did not detect sister cell separation In *Apc^Min/Min^* organoids. Instead, metaphases usually lay in the plane of the epithelium in line with interphase nuclei and only had short compressed basal processes which were difficult to visualise (Figure 6A). In addition, cell morphology was altered and the distance between apical and basal surfaces was significantly reduced (ca. 25%) compared to wild-type or *Apc^Min/+^* cells (Figure 6A). To determine whether this was due to changes in cell shape or overall cell size, we measured the volume of isolated, single cells from wild-type, *Apc^Min/+^* and *Apc^Min/Min^* organoids using flow cytometry. There was no significant difference between wild-type and *Apc^Min/+^* but cell size in *Apc^Min/Min^* organoids was reduced by 25%, indicating that a smaller cell volume was responsible for the reduced cell height (Figure 6B). This suggests that in cells lacking wild-type *Apc*, space restriction causes a reduction in apical-basal distance to prevent INM and restrict basal process formation, preventing post-mitotic separation.

**Figure 6.**
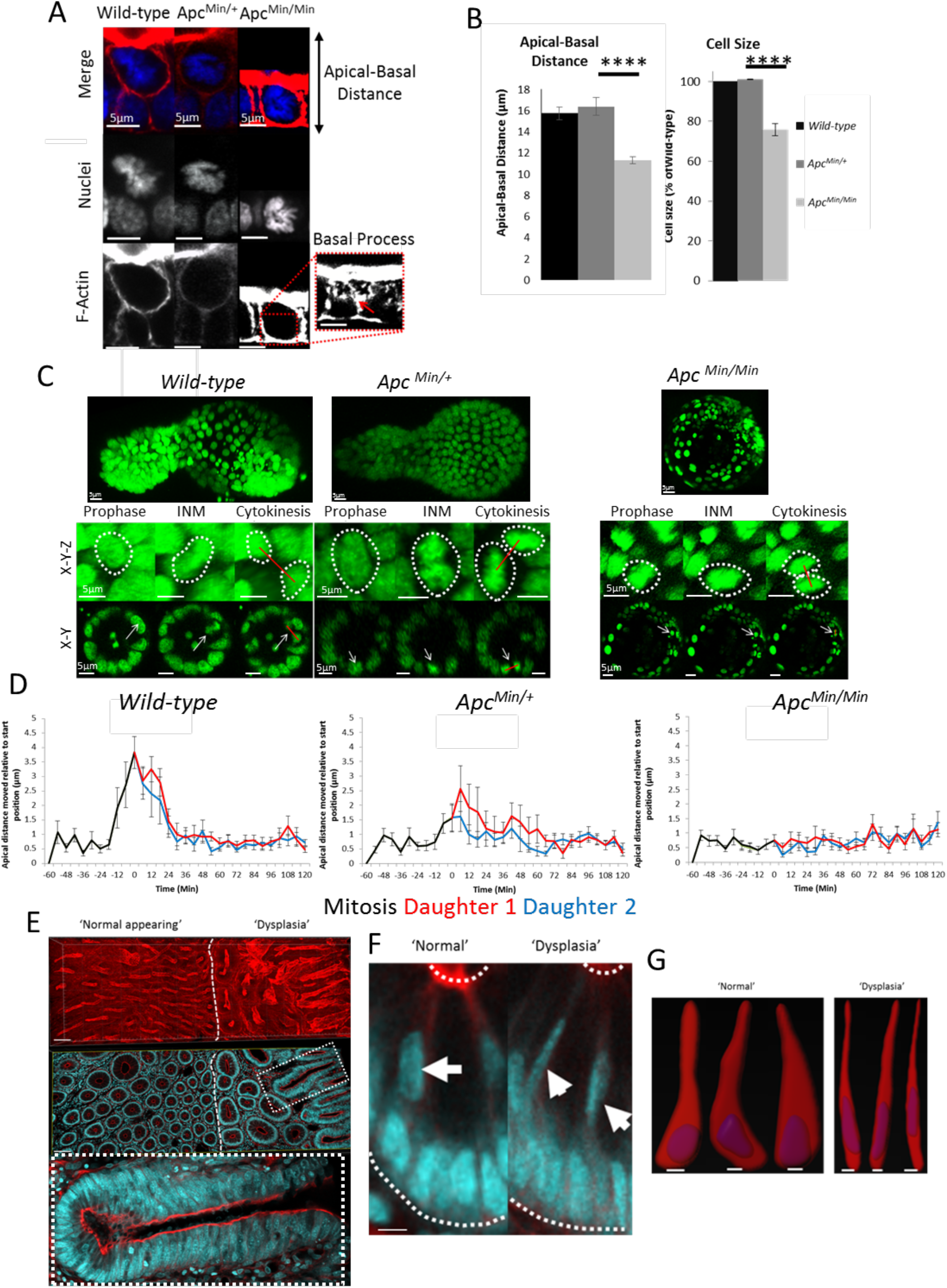
*Apc* mutation limits the ability of daughters to separate by preventing interkinetic nuclear migration and reducing cell size. (**A**) Representative images of metaphases in wild-type, *Apc^Min/+^* and *Apc^Min/Min^* intestinal organoids stained with Hoechst (blue) and phalloidin (red). (**B**) The apical-basal distance of interphase cells was measured in wild-type, *Apc^Min/+^* and *Apc^Min/Min^* organoids in images (left panel). There was a significant difference in the apical-basal distance between wild-type and *Apc^Min/Min^* organoids. Cell size was measured in isolated wild-type, *Apc^Min/+^* and *Apc^Min/Min^* cells using flow cytometry. The median forward scatter was determined from 3 independent organoid samples for each genotype and averaged. Data is displayed relative to the size of wild-type cells. There is a significant difference between the relative cell size of wild-type and Apc^Min/Min^ organoids (T-test, ****p<0.0001) (**C**) Individual frames from H2B-GFP organoid movies show interkinetic nuclear migration (INM). For each genotype, a representative mitosis is shown at prophase, INM and cytokinesis. 3D (maximum intensity projections) and transverse (X-Y) views through an organoid branch or cyst are shown. (**D**) Dynamics of INM during mitosis in wild-type, *Apc^Min/+^* and *Apc^Min/Min^* was measured relative to the starting distance (N = 10 cells per genotype). Data is displayed as mean +/− SEM. Measurements for mother (black line) and daughters (red and blue lines) are superimposed (see also S5 Figure). (**E**) A Vibratome section of human FAP colonic tissue was stained with Hoechst (blue) and phalloidin (red). Displayed are 3D projections (top panel) and section views (bottom panel). Highlighted are regions of ‘normal-appearing’ and ‘dysplastic’ regions. The highlighted area (dashed line) shows a crypt with cell pile-ups/pseudo-stratification (**F**) Magnified view of a crypt ‘normal’ and ‘dysplastic’ regions of FAP colonic tissue in panel (**E**). The basal and apical surfaces are highlighted by white dashed lines. Cells undergoing interkinetic nuclear migration are highlighted by white arrows. Scale bars = 10μm. (**G**) Cell morphology of 3 cells from crypts in ‘normal’ and ‘dysplastic’ regions of FAP colonic tissue. Displayed are surface renders of F-actin and nuclei for individual cells. Scale bars = 5μm.

To directly determine if and how INM was altered in *Apc^Min/Min^* organoids, we first measured the distance of mitotic nuclei relative to the basal membrane of the epithelial layer in wild-type, *Apc^Min/+^* and *Apc^Min/Min^* organoids. The basal reference was established as the plane formed between neighbouring cells proximal to the mitotic/daughter cells (S5 Figure). In wild-type cells, the distance covered by INM was approximately 4μm (Figure 6C, D). There was no significant difference in distance covered during INM between wild-type and *Apc^Min/+^* cells (Figure 6C, D). However, the speed of nuclei during INM was significantly reduced in *Apc^Min/+^* cells suggesting that they require longer to reach the apical region and/or spend less time there (Figure 3D). As expected, in *Apc^Min/Min^* organoids there was no apical displacement and all daughters were placed adjacently. Similar results were achieved in organoids chronically treated with Chir99021 to hyper-activate Wnt signalling (S5 Figure). We also observed mitoses in *Apc^Min/+^* organoids that exhibited no INM. One possible explanation is that some cells in *Apc^Min/+^* organoids had already undergone LOH, which may dramatically reduce INM. Together, these data show that INM is important for the ability of sisters to separate (Figure 6C, D).

To corroborate these observations *in vivo*, we compared the morphology of interphase and mitotic cells in normal and transformed tissue isolated from a familial adenomatous polyposis (FAP) patient (Figure 6E). We detected a striking morphological change in cells from dysplastic regions. In contrast to *Apc^Min/Min^* organoids, which displayed greatly reduced apical-basal distance, we detected significant lateral compression of cells in the human tissue samples (Figure 6F, G) that correlated with the pseudo-stratification caused by ‘pile-ups’ of cells along the crypt-villus axis (Figure 6E). This lateral compression likely restricts the ability of nuclei to undergo INM and reach the apical surface and separate. The resulting decrease in post-mitotic separation may contribute to the observed pseudo-stratification and also promote cell overcrowding in crypts.

## DISCUSSION

Where a cell is born is linked to its identity. In this study, we show that daughter cells can separate immediately after cytokinesis and assume increasingly diverging positions over time (Figure 7). This means that one sister is more likely to exit a compartment where it was born than the other. For stem cells, this means that one sister is more likely to differentiate into a progenitor. For transit-amplifying cells, it means that one sister is more likely to exit the proliferative niche of the transit-amplifying compartment and become post-mitotic. For simple columnar epithelia, it is possible that post-mitotic separation provides a cellular mechanism for the neutral drift that governs stem cell population dynamics. All intestinal cells have a similar probability of undergoing post-mitotic separation, allowing one daughter to remain in its current niche position and the other to leave. It is unlikely that post-mitotic separation always produces a heterogeneous cell pair, as this would only readily occur near a niche boundary. However, this mechanism could influence overall homeostasis and protect stem cell number by slowing neutral drift i.e. ensuring that one daughter remains close to its birth place, making it more likely to remain in a proliferative niche.

**Figure 7.**
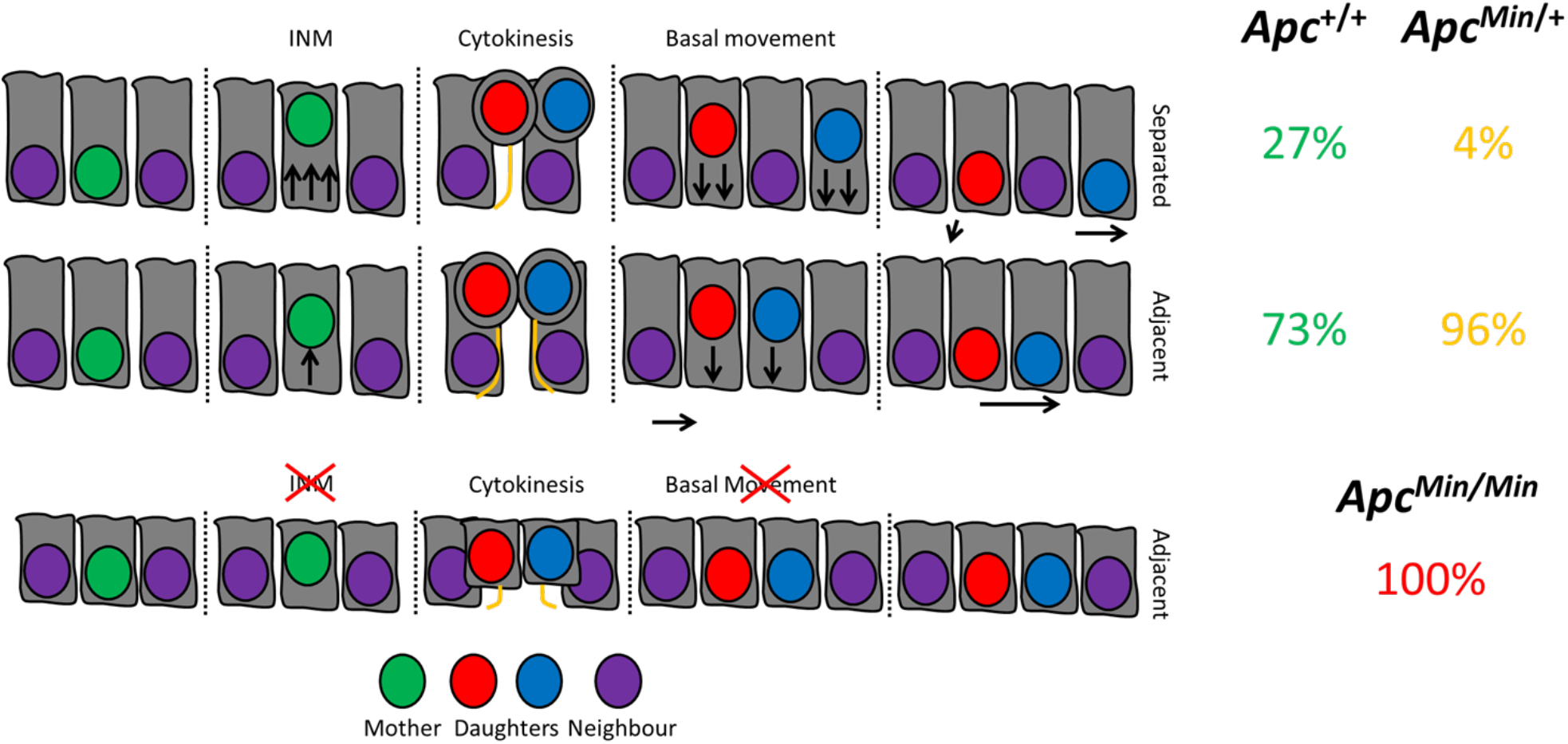
Model for how prolonged niche retention in *Apc* mutant cells arises. In untransformed intestinal epithelia, normal tissue architecture is maintained. During mitosis, cells undergo interkinetic nuclear migration, characterised by the apical movement of nuclei as the cell rounds up. This permits daughter cells to remain proximal or to become displaced from one another by neighbouring cells. Displacement promotes retention of a cell at its birthplace and allows the other to exit the niche. This displacement is facilitated by INM and the asymmetric segregation of the basal process. After mutation of one Apc allele (Apc^Min/+^), mitoses become biased towards adjacent placement and migrate slower, facilitating niche retention. Upon loss of heterozygosity (Apc^Min/Min^), all cells lose the capacity for separation due to reduced cell size and inhibited INM. As a result, symmetrical cyst growth could be promoted, promoting altered tissue architecture.

Reduced post-mitotic separation in *Apc* mutant cells provides an explanation for their increased probability to colonise a niche (Vermeulen et al., 2013, Baker et al., 2014). Neither mutant sister is likely to be displaced from its birthplace, instead, they remain in close proximity to each other. Together with their well characterised decreased migration, which we confirmed in organoids (Figure 4), this could significantly decrease the number of *Apc* mutant cells exiting proliferative compartments (Nelson et al., 2012). As a result, in *Apc* mutant epithelia, many sisters would remain in a proliferative niche, resulting in increased number of proliferating cells. This explains the increased number of cells in the crypt base of *Apc^Min/+^* tissue (Quyn et al., 2010). A reduction in post-mitotic separation and decreased migration may confer on *Apc* mutant cells the competitive advantage that causes their preferred niche retention (Figure 4) (Nelson et al., 2012). Changes in the positioning of wild type and *Apc* mutant cells could also be responsible for the measurable differences in histologically normal appearing Apc^*Min/+*^ tissue. The decrease in the regularity of crypt shape and packing that is detectable by high resolution optical imaging and high frequency ultrasound could reflect altered post-mitotic placement of cells and could be caused by increased retention of expanding clones of *Apc^Min/Min^* mutant cells in *Apc^Min/+^* tissue (Fatehullah et al., 2016a)

Post-mitotic placement is likely to contribute to crypt fission, the process that produces two daughter crypts and is responsible for elongation of intestinal tract (Humphries and Wright, 2008). Initiation of crypt fission involves the formation of a cluster of stem cells at the crypt base, which marks the point of bifurcation (Langlands et al., 2016). Dynamic post-mitotic rearrangements of daughters could explain how these clusters form. We found that in many cases, mitoses next to Paneth cells resulted in separating sisters. The tight packing at the crypt base and the larger size and stiffness of Paneth cells means that once mitotic daughters of a dividing stem cell at the crypt base remain adjacent to each other, it is increasingly difficult for daughters of subsequent divisions to separate due to the physical constraint generated. This could cause the initial clustering of Lgr5+ cells marking the initiation of fission.

Post-mitotic separation is facilitated by INM and the ability to asymmetrically segregate basal processes. A role for APC in INM as suggested by our data is consistent with findings in neuro-epithelia where loss of *Apc* disrupts INM (Ivaniutsin et al., 2009). In the neuro-epithelium, INM relies on microtubules for nuclear movement and actomyosin activity for cell rounding (Spear and Erickson, 2012, Xie et al., 2007). In the intestinal epithelium, INM may also involve microtubules. Specifically, the apical-basal microtubule scaffold may facilitate the nuclear movement during INM (S1 Figure). APC regulates both microtubules and actin (Näthke et al., 1996, Zumbrunn et al., 2001, Okada et al., 2010) and cytoskeletal defects resulting from *Apc* mutation could compromise the function of microtubule bundles reducing the efficiency of INM and sister separation. Indeed, the number of microtubules in large parallel arrays is significantly reduced in *Apc^Min/+^* cells (Mogensen et al., 2002). Disruption of the microtubule scaffold may also cause the defects in cell volume and height observed in Apc^Min/Min^ cells consistent with recent reports suggesting that disruptions of the apical-basal orientation of microtubules can reduce cell height (Toya et al., 2016).

The formation and position of the basal process underlies post-mitotic separation. Unlike previous reports in the colon (Bellis et al., 2012), we demonstrate that asymmetric process localisation actively promotes neighbour exchange and niche exit. How basal processes form is unclear, whether as a cause or a consequence of mitosis. In *Apc* mutant cells, as in the colon (Bellis et al., 2012), processes are usually symmetrically placed and they form more slowly. The increased time required to complete INM in *Apc* mutant cells (Figure 4) may be responsible, by reducing the time available to establish an asymmetric process.

Cell morphology is also important for post-mitotic separation. Cells in highly abnormal regions of FAP tissue were significantly compressed laterally, suggesting that mutant cells are smaller and/or softer than wild-type cells. There is growing evidence that malignant cells are softer than untransformed cells (Plodinec et al., 2012). Reduced cell volume could cause both lateral and/or apical-basal compression and restrict nuclear movement and impair INM. This would cause mutant cells *in vivo* to remain close to their sisters and colonise a niche more successfully than wild-type cells. Altered cell morphology is evident in human intestinal organoids after *Apc* depletion and also seen with mutations in KRas, P53 or SMAD4 (Drost et al., 2015), suggesting that post-mitotic separation can be compromised by other contributing mutations which affect cell morphology.

In summary, we provide evidence that post-mitotic separation is a general mechanism used by intestinal epithelial cells to control niche access. This cellular mechanism could further explain the stochasticity of intestinal homeostasis and how it becomes biased to create a pre-neoplastic state.

## Contributions of authors

T.D.C and I.N designed the study; T.D.C collected the data and performed the analysis; A.J.L assisted with organoid culture and provided images for analysis; J.M.O performed the computational modelling and associated analysis; I.P.N. assisted with animal handling, maintenance and assisted with the scoring of mitotic events; P.L.A. assisted with method development for long-term time-lapse microscopy of organoids; T.D.C and I.N wrote the manuscript with assistance from J.M.O.

## Conflicts of interests

The authors report no conflicts of interest.

## Acknowledgements

We would like to thank members of the Näthke laboratory for general assistance and Dr. Sara-Jane Dunn for helpful discussions. Microscopy and image analysis support was provided by the Dundee Light Microscopy and Tissue Imaging Facility. We would like to also thank Dr. Teemu Miettinen (Univ. of Dundee) for performing flow cytometry to measure cell size; Prof. Mark Chaplain (Univ. of St. Andrews) for helpful discussions on measuring spindle alignment.

## MATERIALS AND METHODS

### Mice

All experiments involving mouse tissue were performed under UK home office guidelines. CL57BL/6 wild-type, Lgr5-EGFP-IRES-creERT2 (*Lgr5^GFP/+^*), *Apc^Min/+^* and R26-rtTA Col1A1-H2B-GFP (H2B-GFP) mice were sacrificed by cervical dislocation or CO_2_ asphyxiation.

### Tissue Preparation: Mouse Small Intestine

Adult mouse small-intestine was washed briefly in PBS and fixed with 4% PFA for 3 hours at 4°C. Intestine was cut into 2x2cm^2^ pieces and fixed in 4% PFA overnight at 4°C. The tissue was embedded in 3% low melting temperature agarose and sectioned at 200μm intervals using a Vibratome (Leica). Cut sections were washed in PBS and permeabilized for 2 hours with 2% Triton X-100 and incubated with Blocking Buffer (1% BSA, 3% Normal Goat Serum, 0.2% Triton X-100 in PBS) for 2 hours at 4°C. Tissue was incubated for 48 hours with Hoechst 33342 (Thermo Fisher, 1:500) and phalloidin (Molecular Probes, 1:150) diluted in Working Buffer (0.1% BSA, 0.3% Normal Goat Serum, 0.2% Triton X-100 in PBS) at 4°C. The tissue was washed with PBS before mounting in Prolong Gold. Sections were mounted on coverslips between 2x120μm spacers to preserve tissue structures.

### Organoid culture

Organoids were generated from mouse small intestinal crypts as previously described (Sato et al., 2009). Briefly, small intestines were removed and washed in PBS and opened longitudinally. Villi were removed by scraping the lumenal surface with a coverslip. Tissue was washed in PBS, incubated in 30mM EDTA (20 minutes) and crypts dislodged by vigorous shaking. Crypt suspensions were centrifuged (600rpm, 4°C) and the pellet washed twice in PBS and dissociated to single cells with TripLE Express (Life Technologies) at 37°C for 5 mins. Cells were resuspended in Advanced DMEM/F12 (ADF) and filtered through a 40μm cell strainer (Greiner). Single cells were resuspended in growth factor reduced, phenol red-free Matrigel (BD Biosciences). Organoids were grown in crypt media (ADF supplemented with 10mM HEPES, 2mM Glutamax, 1mM N-Acetylcysteine, N2 (Gemini), B27 (Life Technologies), Penicillin-Steptomycin (Sigma-Aldrich), growth factors (EGF, 50ng/ml; Invitrogen, Noggin (100ng/ml; eBioscience), and R-Spondin conditioned media (1:4). Chiron99021 (3μM; Invitrogen), valproic acid (1mM; Invitrogen) and Y27632 (10μM; Cambridge Bioscience) were added to organoids for the first 48 hours. Organoids were passaged by physically breaking up Matrigel, washing in ADF, dissociation by pipetting and reseeding in Matrigel.

### Human tissue

Human tissue used in this study was the same as used for a previous study (Fatehullah et al., 2016b). All tissue collected was approved by the Tayside Tissuebank subcommittee of the Local Research Ethics Committee and obtained in accordance with approved guidelines. FAP biopsies from one FAP patient was obtained during routine colonoscopy surveillance.

### Organoid Immunofluorescence

Organoids were grown in 8-well chamber slides (Ibidi) for 1-2 days at 37°C, 5% CO_2_. Organoids were fixed with warmed 4% PFA in PBS (pH 7.4) for 30 minutes (37°C), permeabilized for 1 hour in 1% Triton X-100 (this and subsequent steps were performed at room temperature), blocked for 1 hour (1% BSA, 3% Normal Goat Serum, 0.2% Triton X-100 in PBS). Organoids were incubated overnight in primary antibodies diluted in Working Buffer (0.1% BSA, 0.3% Normal Goat Serum, 0.2% Triton X-100): g-tubulin (Sigma: T6557, 1:500), GFP (Abcam: ab13970, 1:500); β4-Integtrin (Abcam: ab25254, 1:100); YL1/2 (1:200), washed 5x with Working Buffer before overnight incubation with secondary antibodies diluted in Working buffer: Alexafluor™ conjugated (1:500, Molecular Probes) along with 5μg/ml Hoechst 33342 and Alexafluor™ conjugated phalloidin (1:150). Organoids were mounted in Prolong Gold overnight.

### Microscopy

Images of tissue and organoids were acquired with a Zeiss LSM 710 or LSM 880 with Airyscan (Carl Zeiss) using 25X or 40X Zeiss objective lenses and immersion oil with refractive index of 1.514. Serial image stacks were acquired with an optical section size of 0.8μm.

### Confocal Live Imaging

Organoids were grown in Matrigel and spread thinly onto 35mm^2^ glass bottom dishes (World Precision Instruments). Crypt media was supplemented with 2mg/ml doxycycline to induce H2B-GFP expression. For live-cell imaging of mitochondrial dynamics, induced H2B-GFP organoids were incubated with 500nM Mitotracker DeepRed FM (ThermoFisher Scientific) in crypt media for 1 hour; 37°C 5% CO_2_. Subsequently, staining media was replaced with fresh crypt media containing growth factors. For live-imaging of actin, organoids were stained with 100nM SiR-Actin in crypt media overnight. Organoids were placed in a live-cell imaging chamber attached to a Zeiss 710 confocal microscope maintained at 37°C, 5% CO_2_. Images were acquired using a 40X Zeiss immersion objective. Serial image stacks were acquired at optimal interval sizes using minimal laser power every 6 minutes.

### Spindle Angle measurements

Spindle orientation was measured using image stacks and analysed using the Imaris imaging software (Bitplane). Surfaces of the Hoechst, g-tubulin and phalloidin signals were rendered using the isosurface tool. Similar to a report in the mouse colon (Bellis et al., 2012), we used two angles to represent spindle orientation: 1) relative to the crypt axis (Axial angle) and 2) relative to the apical surface (Apical angle). The apical surface was defined using the F-actin signal at the lumenal surface. To calculate angles, three sets of measurement points were manually placed in 3D:

(1) Two points defining the two centrosomes. The connection between them represents the spindle axis.
(2) Two points placed at either end of the crypt so that the axis formed between them is representative of the crypt axis.
(3) Three points placed on the rendered phalloidin surface to represent the cells apical surface.

Axial angles are the angles between the spindle and crypt-axis. The Apical angle is calculated using the 3 measurement points to determine the normal surface vector to the apical plane. The Axial angle between this defined apical surface and the spindle axis was calculated as follows:

#### Calculating the Axial angle

The Spindle-axis vector 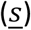 is calculated using Centrosome point 1, [*S*_*x*1_, *S*_*y*1_, *S*_*z*1_], and centrosome point 2, [*S*_*x*2_, *S*_*y*2_, *S*_*z*2_], (**Equation 1**). The crypt-axis vector 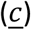 is calculated using crypt-axis point 1, [*C*_*x*1_, *C*_*y*1_, *C*_*z*1_], and Crypt-axis point 2, [*C*_*x*2_, *C*_*y*2_, *C*_*z*2_], (**Equation 2**).

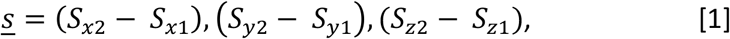

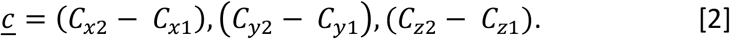

The *α*-angle is calculated by projecting the Vector 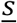 on Vector 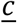 (**Equation 3**)

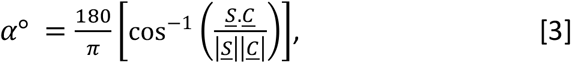

where

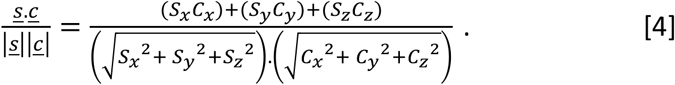

##### Calculating the Apical angle

The Apical angle is calculated using three apical surface points (AP): [*A_x_, A_y_, A_z_*], [*B_x_, B_y_, B_z_*] and [*C_x_, C_y_, C_z_*], placed on the plane, which approximates the apical surface. The coordinates of these points are used to determine two vectors, 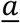 and *ḇ* (**Equation 5, 6**).

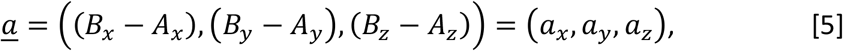

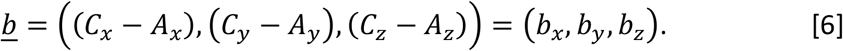

These vectors can subsequently be used to determine the normal surface vector 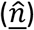 to the apical plane by finding the cross product between vectors 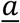 and *ḇ*. (**Equation 7**).

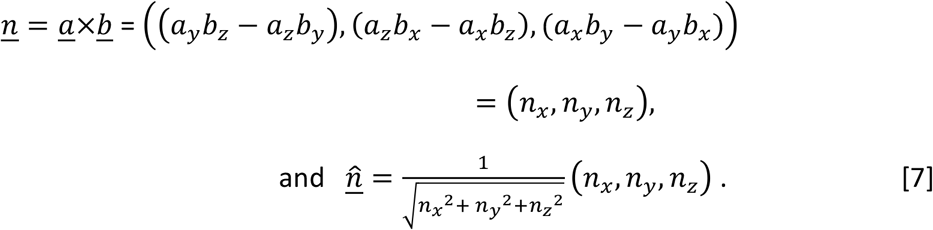

The normal surface vector can then be used to determine the angle between the spindle vector 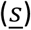 and the surface plane (**Equation 7**).

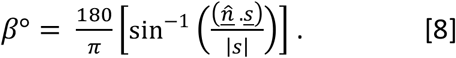

Given that 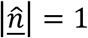.

### INM Measurements

Apical and basal interkinetic nuclear migration was measured by determining the distance between the nucleus and the plane of the epithelium. The plane of the epithelium is defined as the plane formed between the neighbours of the query nucleus:

Find the absolute distance of the mitotic cell, *M*(*x*_0_, *y*_0_, *z*_0_) to epithelial plane, *P*(*Ax* + *By* + *Cz* + *D* = 0). Where the epithelial plane is defined by the plane formed between 3 neighbouring epithelial cells.

Distance to the epithelial plane 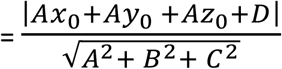

Distance measurements were calculated for 10 planes encompassing each permutation of neighbouring cells. The average distance for these 10 planes was taken as the representative distance of the query cell in reference to the plane of the epithelium in which it originated. This was to account for variability induced by the curvature within the organoid branches.

### Definition of Stem Cell and Transit Amplifying compartments

Stem cell and transit amplifying compartments were defined based on the average position of Lgr5+ cells along the crypt-villus axis measured in Lgr5-GFP organoids (S5 Figure). As Lgr5 can also be expressed in the early TA compartment (Quyn et al., 2010), we conservatively defined the SC compartment based on the average position of Lgr5(+) cells rather than the average distance of the Lgr5(+) cell furthest from the crypt base.

### Distance measurements

Cell position was determined by placing a measurement point in the centre of each nucleus. The co-ordinates of individual points were used to measure the distances between nuclei or from the crypt base. The crypt base was defined by a reference nucleus manually chosen at the crypt base. The reference nucleus was determined at each time point. Distance between points was calculated by the standard formula:

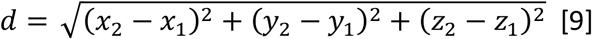

### Time-lapse analysis and scoring

Time-lapse image stacks were analysed manually using Imaris (Bitplane). All mitotic events were marked using the ‘3D spots function’. Time-point 0 was denoted as the time-point immediately before cytokinesis. Each mitotic daughter cell was tracked until its death, subsequent division, or exit from the imaging window. If daughter cells were separated by a neighbouring nucleus after basal INM (at 120 minutes) they were scored as separated. Displacement from the ‘point of birth’ was calculated as the change in distance between the start (mitotic mother) and end position of a daughter, divided by the time interval. Daughter 1 was always classified as the daughter closest to the base of the crypt.

### INM measurement

We determined the distance covered by nuclei during INM by measuring the distance of the mitotic nucleus to a defined basal reference plane. The basal reference was defined as the plane formed by neighbouring interphase cells. The co-ordinates of 5 neighbouring nuclei most proximal to the mitotic cell were determined. These interphase cells often did not form a simple plane due to curvature of the epithelial sheet. To account for the shape, we calculated the nearest distance from the position of the mitotic cell to the planes formed by combinations of each of three nuclei for the 5 neighbours. The average distance from these (10) planes was then used as the estimated apical distance travelled during INM (S5 Figure). Please refer to supplementary information for further details.

### Cell height measurement

Apical-basal distance was measured as the distance between the apical and basal surfaces using Imaris. Distances were calculated in the optical section at the centre of the chosen cell by recording one 3D measurement point in the middle of the apical and one in the middle of the basal surface. The distance between these points was the apical-basal distance.

### Sample size and Statistical analysis

Analyses were performed using at least three different organoids. Individual cell comparisons were performed using at least 10 cells. Comparisons had to be made between organoids imaged in separate imaging sessions because each time lapse took three days. All statistical tests were performed using Prism 6.0a (GraphPad). Tests were performed as described in figure legends (significance: ns = not significant, *p<0.05, **p<0.01, ***p<0.001, ****p<0.0001).

### Multicellular Computational Model

All simulations were undertaken in the CHASTE framework (Mirams et al., 2013). We extended the model presented in (Dunn et al., 2016) to permit variable separation of cells after division. In summary, cells are represented by their centres, which are free to move on a surface (in 3 dimensional space) that is defined using measured crypt geometry. Cells move because of forces exerted on them due to compression of, and by, neighbouring cells. We are using the optimal model identified in Dunn et al., 2016, Model 6. In this model cells divide after a uniformly distributed time that depends on the level of Wnt (imposed as a linear gradient) experienced when the parent cell divided. Additionally, if cells are compressed beyond a given threshold they pause in the G1 phase of the cell cycle. All parameters used are as described in Dunn et al., 2016. As in previous three-dimensional models of the crypt (Dunn et al., 2016, Dunn et al., 2013), cell division occurs in a direction uniformly drawn from the sphere surrounding the centre of the dividing cell. Daughter cells are placed at a specified distance from one another. Previously, in all existing models, this distance is chosen so daughter cells are adjacent to each other. We modified this parameter so that two thirds of all cell divisions resulted in the daughter cells being placed next to each other (a separation of 1 cell diameter), the remaining third resulted in the daughter cells being separated by *S* cell diameters. We vary this parameter between 0 and 3 to measure the effects on separation of cells in the virtual crypt.

## Supplemental Information

We provide supplemental information describing mitosis in intestinal epithelium and organoids, showing that they are similar and suggest that INM requires microtubules (S1 Figure). In addition to asymmetric basal tethering, post-mitotic separation appears to be influenced by the proximity of other mitotic daughters and by proximity to Paneth cells which are less mobile (S2 Figure). We also show that asymmetric tethering and spindle orientation are partially linked and that this is altered in *Apc* mutant tissue (S3 Figure). We show that disruption of INM can also be induced by hyper-activation of Wnt (S4 Figure). We provide figures demonstrating our definition of the stem and transit amplifying compartments and an illustration of how INM was measured (S5 Figure).

### Movie Legends

**S1 Movie. H2B-GFP Intestinal Organoids**

Confocal LSM imaging of induced H2B-GFP organoids derived from wild-type and Apc^Min/+^ mice (Both untransformed (Apc^Min/+^) and transformed cysts (Apc^Min/Min^)).

**S2 Movie. Adjacent Sister Placement**

Confocal LSM imaging of an induced wild-type H2B-GFP organoid showing manual tracking of a mitotic cell and its progeny. Daughters were tracked manually using Imaris. In this example, both daughter cells ‘re-insert’ into the epithelium as neighbours.

**S3 Movie. Post-mitotic Separation**

Confocal LSM imaging of an induced wild-type H2B-GFP organoid showing manual tracking of a mitotic cell and its progeny. Daughters were tracked manually using Imaris. In this example, the ‘blue’ daughter cell is displaced from its sister.

**S4 Movie. 4D Visualisation of post-mitotic separation**

Confocal LSM imaging of an induced wild-type H2B-GFP organoid showing manual tracking of a mitotic cell and its progeny undergoing post-mitotic separation. Surface rendering was performed to highlight the mother (cyan), sisters (blue/red) and neighbour cells (magenta). The respective timelapse is shown in the top panels and a 3D rotation around the timepoints encompassing interphase, INM, cytokinesis and after separation are displayed in the bottom panels.

**S5 Movie. Mitotracker in Intestinal Organoids**

Confocal LSM imaging of an induced H2B-GFP organoid treated with Mitotracker.

**S6 Movie – Mitotic neighbours**

Confocal LSM imaging of an induced wild-type H2B-GFP organoid showing manual tracking of a mitotic cell and its progeny undergoing post-mitotic separation. In this example, the daughters of the original mitosis (red) are displaced by the placement of a daughter cell from an adjacent mitosis (purple).

**S7 Movie – Paneth Cell proximity and displacement**

Confocal LSM imaging of an induced wild-type H2B-GFP organoid showing manual tracking of a mitotic cell and its progeny undergoing post-mitotic separation. In this example, the daughters of this mitosis are displaced in proximity to a Paneth cell (recognisable by the large space with no nuclei). This movie was used in stills in S2 figure.

**S8 Movie – Delayed Daughter Cell Insertion**

Confocal LSM imaging of an induced wild-type H2B-GFP organoid showing manual tracking of a mitotic cell and its progeny undergoing post-mitotic separation. In this example the left most daughter takes two attempts to reassume its interphase position, whilst the other is displaced. This movie was used for stills in Figure 5E.

**S9 Movie – Asymmetric process segregation underlies post-mitotic separation**

Confocal LSM imaging of an induced wild-type H2B-GFP organoid treated with SiR-Actin. The movie shows a mitotic cell undergoing post-mitotic separation in which one daughter retains the basal process. The two daughters (white spheres) are separated by a neighbour after reinsertion.

## Supplemental Figure Legends

**S1 Figure.**
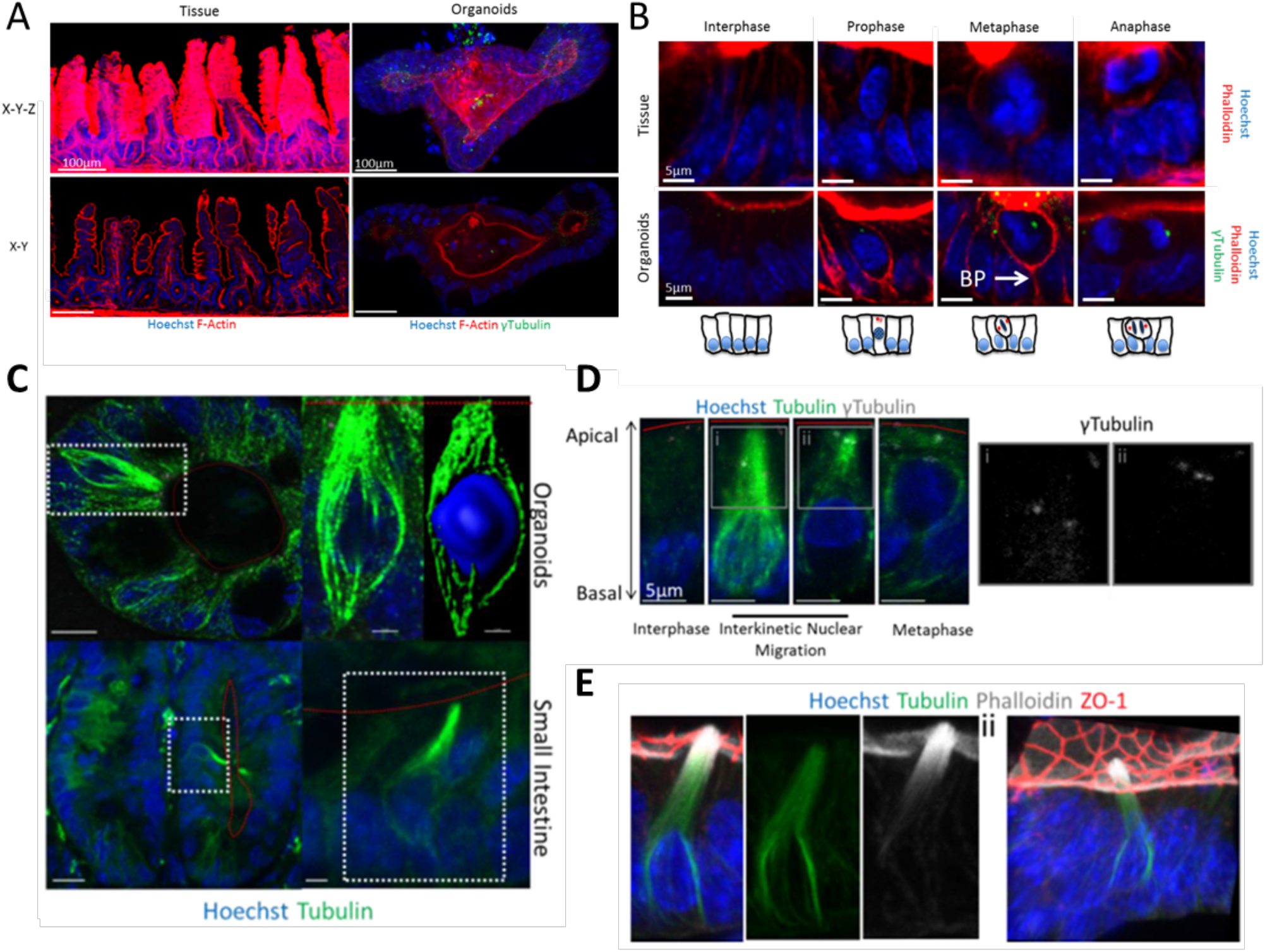
Mitosis in intestinal tissue and intestinal organoids. (**A**) Maximum intensity projection (X-Y-Z) and confocal sections (X-Y) of a vibratome section of mouse small-intestine (tissue) and an intestinal organoid stained with Hoechst (blue), phalloidin (red), and γ-tubulin (green). (**B**) Confocal sections (X-Y) of mitotic stages visualised in intestinal crypts of whole tissue and organoids. Interphase cells maintain basally positioned nuclei. During mitosis the apical cell surface remains aligned with neighbouring cells. Chromatin condensation occurs during prophase and the nucleus is displaced apically. During INM, the rounded mitotic cell remains attached to the basal membrane by a basal process (BP). After alignment with the apical surface the metaphase plate forms apically and is directly followed by anaphase in which cells have two clear sets of sister chromatids. (**C**) An intestinal organoid and small intestinal tissue stained with Hoechst (blue) and an antibody against tubulin (green). In the right panel, surface rendering reveals the structure of the apical-basal array of microtubules. The apical surface is marked by the red dashed line. (**D**) Representative examples of mitotic cells in organoid epithelium at stages of interkinetic nuclear migration. Microtubule polymerisation is most evident when nuclei are basally localised. As nuclei move apically, microtubule polymerisation is mostly at the apical-most side of the microtubule scaffold. There was no detectable microtubule polymerisation in rounded up mitotic cells. Indicated cells are not differentiated due to the detectable pair of centrosomes (i and ii). The apical surface is marked by the red dashed line. (**E**) A tuft cell in an intestinal organoid. Organoids were stained with Hoechst (blue), phalloidin (white) and antibodies against tubulin (green) and ZO-1 (red) displayed as a section (i) or in 3D (ii). Tuft cells have a ‘tuft’ of microvilli that protrude apically into the lumen. They are fully differentiated, predominantly found within the differentiated zone, and lack detectable centrosomes.

**S2 Figure.**
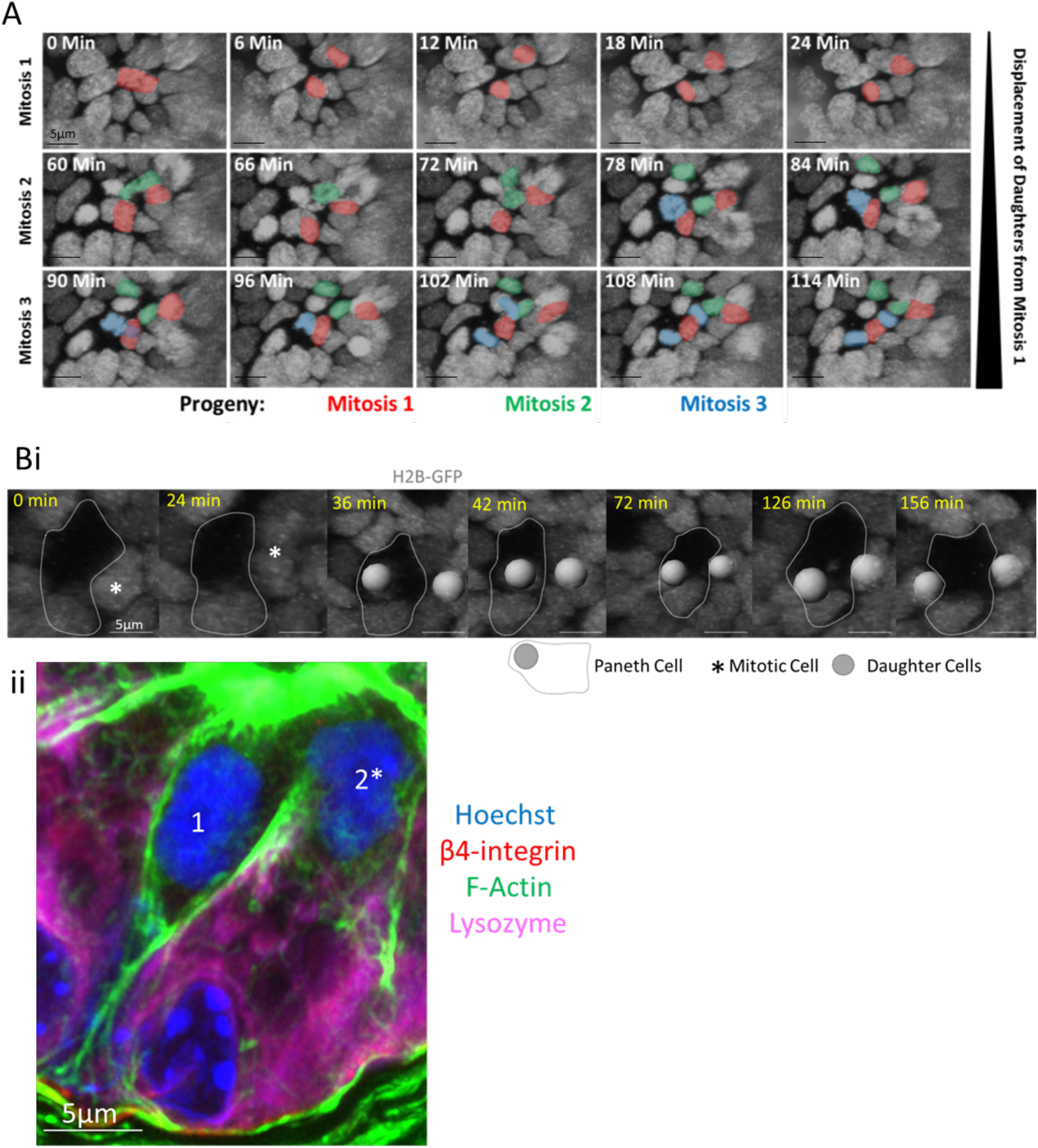
Alternative methods of separation. (**A**) Separation of daughters is enhanced by movement of neighbouring mitotic cells. Displayed are 3D projections of the movements of the progeny of a mitotic cell (Original Mitosis, [Mitosis 1; red]) and the progeny of two neighbouring mitotic cells (Mitosis 2, green; Mitosis 3; blue). Time 0 marks metaphase of the original mitotic cell. (**B**) **i**)Live imaging of a wild-type H2B-GFP organoid. A Paneth cell can be clearly identified based on morphology and distribution of neighbouring nuclei (dashed line). A mitotic cell (white stars) proximal to the Paneth cell divides to produce two daughters (white balls) who separate and then renter the epithelial plane adjacent to the Paneth cell. ii) Fixed image of small-intestinal tissue, stained with Hoechst (Blue), β4-Integrin (red), lysozyme (magenta) and phalloidin (green). A recent mitosis produced two daughter cells (1, 2*) which have become separated and have reinserted on either side of the Paneth cell. The image is the same as that in Figure 5A, showing lysozyme staining.

**S3 Figure.**
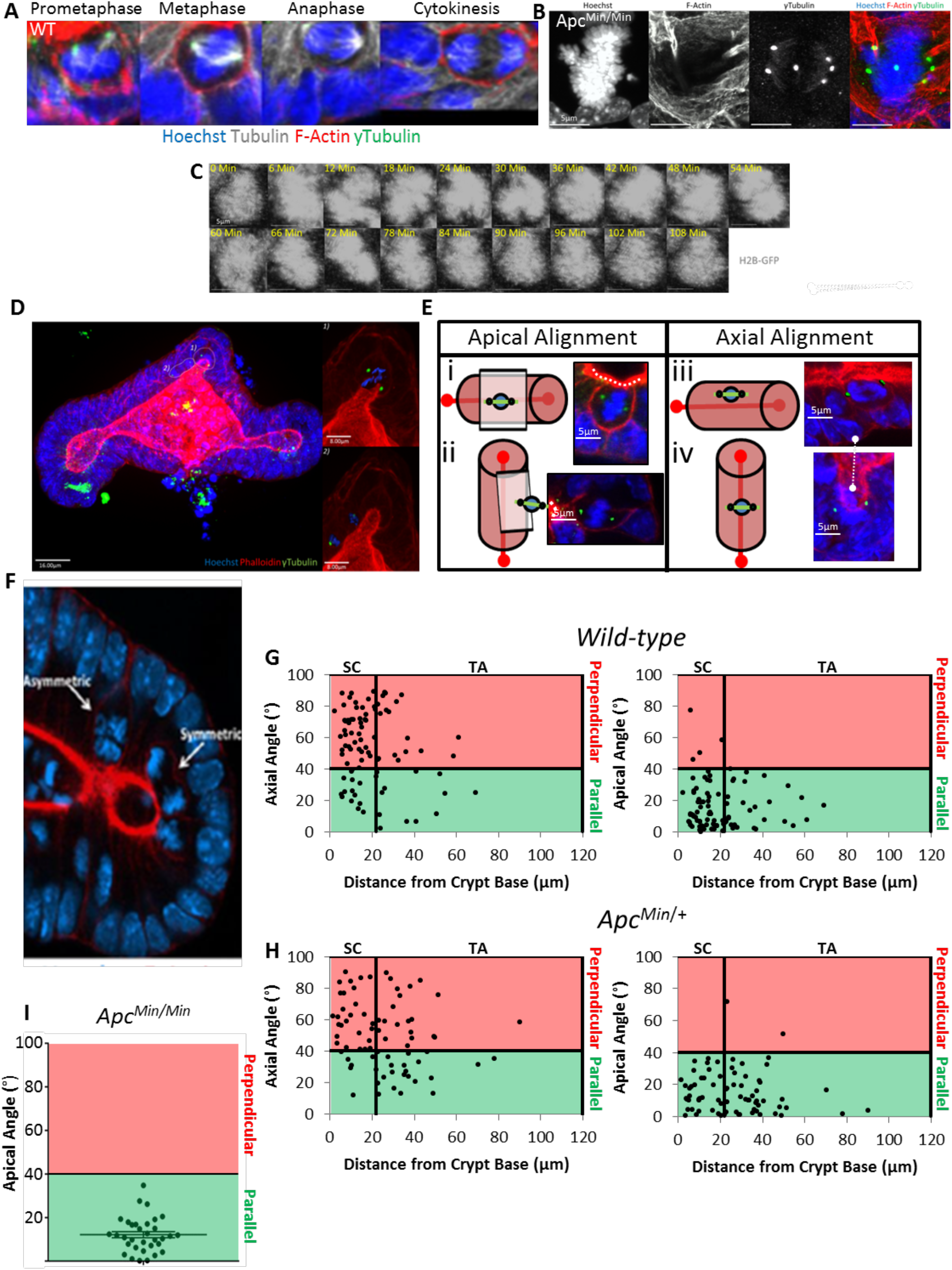
Spindle orientation in intestinal organoids. A) Representative mitotic cells in prometaphase, metaphase, anaphase and during cytokinesis. Organoids are stained with Hoechst, phalloidin and antibodies against tubulin and y-tubulin to visualise nuclei (blue), F-actin (red), microtubules (white) and centrosomes (green). Centrosomes are located equidistantly on either side of the metaphase plate once it is fully established. B) A representative example of an Apc^Min/Min^ mitotic cell with a multipolar spindle. Apc^Min/Min^ organoids were stained with Hoechst (blue), phalloidin (red) and an antibody against γ- tubulin (green) to visualise DNA, F-actin and centrosomes. C) A representative example of an Apc^Min/Min^ cell undergoing mitotic slippage. Displayed are stills from live-imaging of an Apc^Min/Min^ H2B-GFP organoid. Chromosome condensation is clearly observed as the cell enters prophase. Instead of proceeding with mitosis, chromosomes de-condense as the cell returns to interphase. D) A representative example of a wild-type organoid stained with Hoechst (blue), phalloidin (red), γ-tubulin (green) to visualise DNA, F-actin and centrosomes. Two mitotic cells are highlighted, one in metaphase (top) and one in anaphase (bottom). Surface rendering in Imaris can clearly highlight individual cells and their two centrosomes. E) Potential spindle alignments in intestinal organoids. Spindle orientation was determined in reference to: 1) the axis of tissue growth; the crypt-villus axis or 2) the apical surface. Spindle orientations are the angle between the spindle and crypt-villus axis (Axial angle) or apical surface (Apical angle). Examples of each type of spindle alignment is displayed; i) Parallel to the crypt-villus axis (Crypt lengthening), ii) Perpendicular to the crypt-villus axis (Crypt widening), iii) Parallel to the apical surface (‘symmetric’ division) or perpendicular to the apical surface (‘asymmetric’ division). Reference axes are highlighted by the white dashed line. F) Representative example of an asymmetrically and symmetrically oriented division in the crypt base of an intestinal organoid. The organoid is stained with Hoechst (blue) and phalloidin (red). Note that a pro-daughter cell in the asymmetrically aligned mitoses is poised to inherit the basal process. Spindle orientations were determined for mitoses in G) wild-type, H) *Apc^Min/+^* and I) *Apc^Mm/Min^* organoids. Only apical angles could be calculated for Apc^Min/Min^ organoids due to loss of crypt-villus architecture. Data is displayed in reference to the crypt base. Angles greater than 40° were classified as perpendicular. Angles less than 40° are classified as parallel. Data is displayed as a function of distance along the crypt-villus axis. The stem cell compartment was defined as the curved region at the base of branches, approximately 20μm from the lumenal crypt base.

**S4 Figure.**
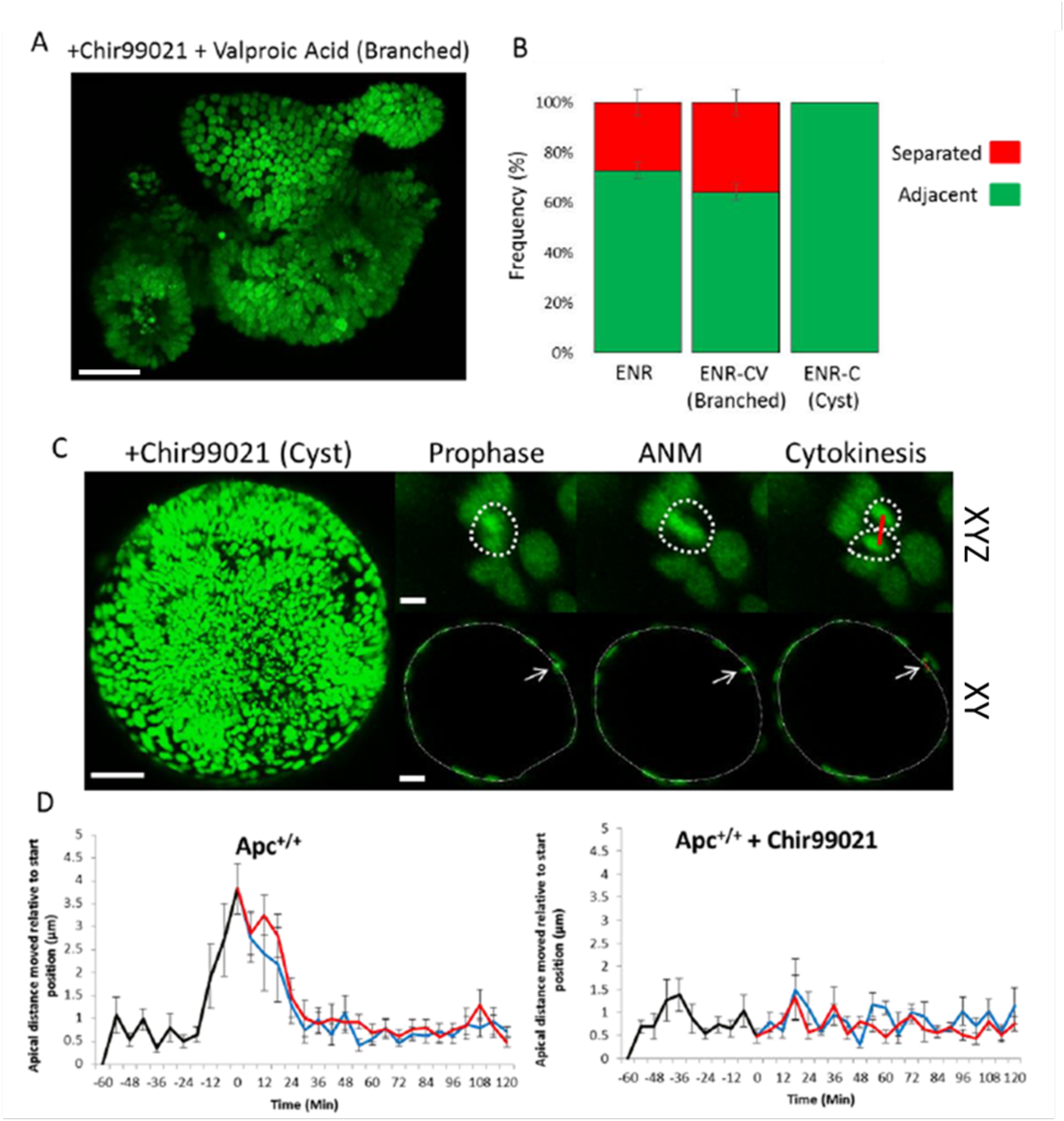
Disruption of INM can be induced by chronic Chir99021 treatment. **A**) An H2B-GFP intestinal organoid treated with Chir99021 and valproic acid. Treatment with Chir99021 and valproic acid has been shown to increase the frequency and distribution of Lgr5(+) cells along the crypt axis (Yin et al., 2014). Treated organoids retain crypt-villus architecture. Scale bar = 100μm **B**) Daughter cell placement after mitosis was scored in H2B-GFP organoids using time-lapse movies. Organoids were treated with Chir99021 and valproic acid to increase the stem cell content along the crypt-villus axis, or they were chronically treated with 10μM Chir99021 for 4 days to induce cyst formation in WT organoids. Division subtypes were compared to untreated organoids (ENR). Data was compared to the dataset in Figure 4B. (ENR N = 6 organoids, N = 491 mitoses; ENR-CV (branched) N = 3 organoids, N = 351 mitoses; ENR-C (cyst) N = 3 organoids). **C**) A wild-type H2B-GFP organoid chronically treated with 10μM Chir99021 for 4 days (left panel), scale bar = 100μm. Right panels highlight individual frames from live-recordings displaying a representative mitosis during prophase, apical interkinetic nuclear migration (ANM) and cytokinesis. 3D (maximum intensity projections, X-Y-Z) and transverse (X-Y) are shown. Scale bars = 5μm **D**) Dynamics of interkinetic nuclear migration during mitosis in Chir99021 treated organoids *were* measured relative to the starting distance (N = 10 cells). Data is displayed as mean +/− SEM. Measurements of the mother (black line) and daughter cells (red and blue lines) are superimposed. The wild-type dataset displayed is the same dataset as displayed in Figure 7D.

**S5 Figure.**
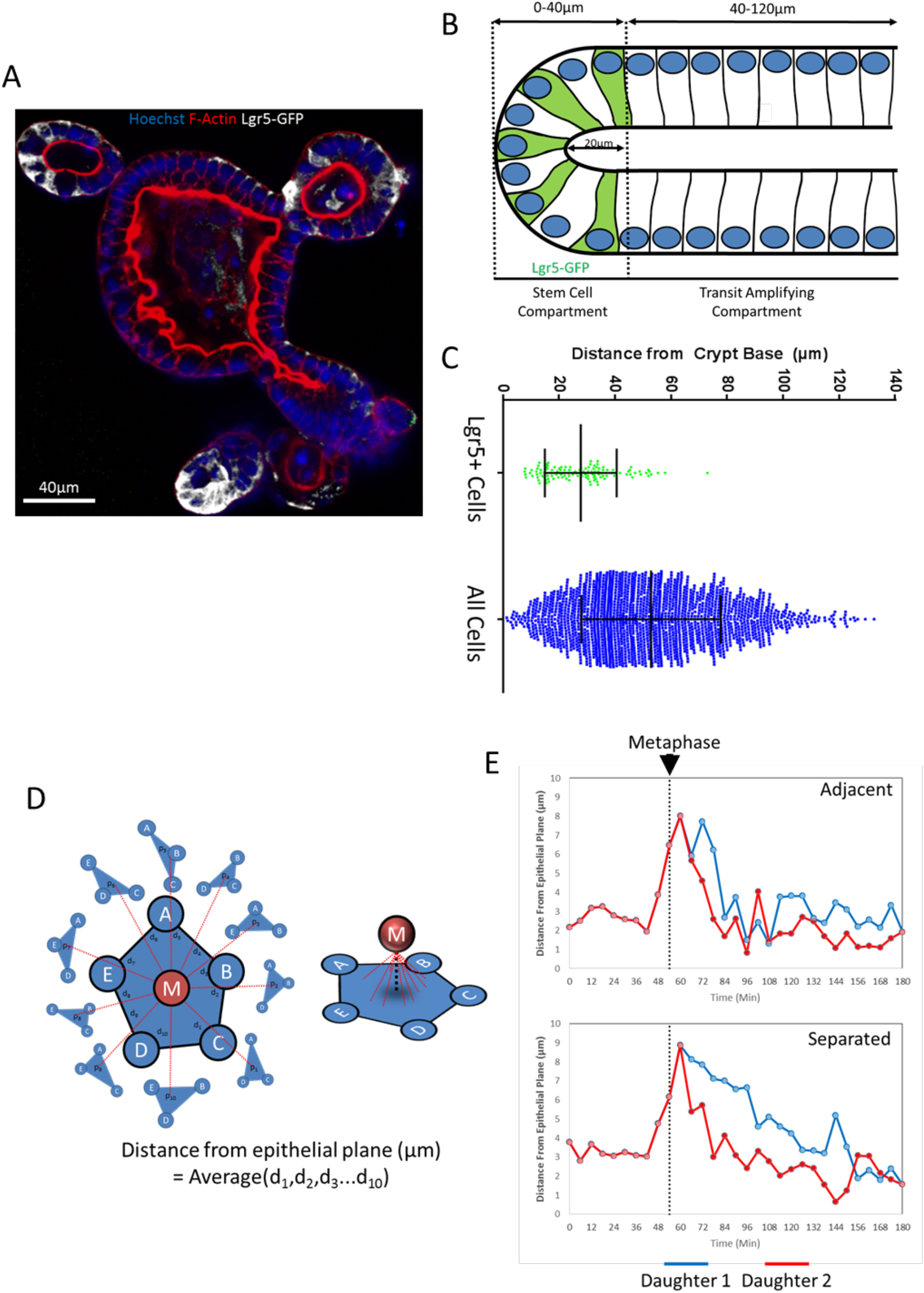
Definition of tissue compartments and interkinetic nuclear migration. **A**) An Lgr5-GFP expressing intestinal organoid stained with Hoechst (nuclei), phalloidin (F-actin) and GFP (Lgr5+ stem cells). Stem cells resided within the base of intestinal organoid branches, mostly residing within the curved region of the crypt base. **B**) The position of each Lgr5-GFP+ cell was recorded and compared to the positions of the total cell population with reference to the crypt base. Nuclear position was used as a surrogate for cell position and distances were compared to the nucleus closest to the base of the crypt. Data was pooled from 6 individual organoids. Data is displayed as mean +/− SD. **C**) Diagram showing the defined compartments within intestinal crypts. The majority of Lgr5+ stem cells were located approximately 0-40μm from the crypt base. This region was termed the stem cell compartment. This equated to the curved region at the base of the crypt, approximately 20μm from the lumenal crypt base. Above this region we defined as the transit-amplifying compartment. A small fraction of GFP+ cells resided above the defined stem cell compartment, similar to our previous studies in whole intestinal tissue. **D**) Interkinetic nuclear migration is quantified as the distance of the query cell in reference to the epithelial plane in which it originated. The plane of the epithelium is defined as the plane in which neighbouring nuclei are located. A plane is defined by the co-ordinates of 3 points. Therefore the distance was measured between the query cell and the plane formed by 3 of its neighbour nuclei. This process was repeated utilizing 5 neighbour cells. The average distance for each of these 10 planes was determined as distance from the epithelial plane. **E**) Examples of INM measurement for a cell undergoing adjacent placement (Adjacent) or post-mitotic separation (Separated). Distances were determined for each time-point during prophase and for each of the daughter cells (red and blue lines).

